# A multimodal approach to identify clinically relevant parameters to monitor disease progression in a preclinical model of neuropediatric disease

**DOI:** 10.1101/522011

**Authors:** Tyler B. Johnson, Jon J. Brudvig, Kimmo K. Lehtimäki, Jacob T. Cain, Katherine A. White, Timo Bragge, Jussi Rytkönen, Tuulia Huhtala, Derek Timm, Maria Vihma, Jukka T. Puoliväli, Antti Nurmi, Jill M. Weimer

## Abstract

While research has accelerated the development of new treatments for pediatric neurodegenerative disorders, the ability to demonstrate the long-term efficacy of these therapies has been hindered by the lack of convincing, noninvasive methods for tracking disease progression both in animal models and in human clinical trials. Here, we unveil a new translational platform for tracking disease progression in an animal model of a pediatric neurodegenerative disorder, CLN6-Batten disease. Instead of looking at a handful of parameters or a single “needle in a haystack”, we embrace the idea that disease progression, in mice and patients alike, is a diverse phenomenon best characterized by a combination of relevant biomarkers. Thus, we employed a multi-modal quantitative approach where 144 parameters were longitudinally monitored to allow for individual variability. We use a range of noninvasive neuroimaging modalities and kinematic gait analysis, all methods that parallel those commonly used in the clinic, followed by a powerful statistical platform to identify key progressive anatomical and metabolic changes that correlate strongly with the progression of pathological and behavioral deficits. This innovative, highly sensitive platform can be used as a powerful tool for preclinical studies on neurodegenerative diseases, and provides proof-of-principle for use as a potentially translatable tool for clinicians in the future.

**One Sentence Summary:** Principal component analysis identifies a set of clinically relevant parameters able to measure progression of Batten disease in a mouse model.

## Introduction

Rare diseases, conditions that affect fewer than 200,000 patients in the U.S. or less than 1 in 2,000 people in the EU (*1*), represent a particular challenge for medical diagnosis as clinical features are often complex and enigmatic. While very few rare diseases have effective treatments, resulting from the limited information that is typically available for many of these conditions, access to improved animal models and state-of-the art medical diagnostic capabilities are helping to accelerate the number of clinical trials and treatments available to patients. Due to their rarity, access to patients is particularly limited, so researchers and clinicians must rely on comprehensive natural history studies that provide a snapshot of where a typical patient would exist in time. Moreover, because many of these diseases are pediatric and ultimately fatal, the Rare Diseases Act of 2002 made it possible to accelerate the clinical trial design process, with one concession being that the trial need not include untreated controls, thus making the natural history data even more essential. Although much attention has been focused on collecting this information, much of what is captured can be subjective and qualitative. Thus, quantitative biomarkers that can be monitored longitudinally and are minimally invasive are greatly needed in order to monitor treatment responses in both preclinical animal models and human clinical trials. Translational utilization of animal models of human disease benefits greatly from relevant phenotypic characterization. Unfortunately, the techniques most commonly used in animal models often suffer from a lack of translatability. Behavioral assays for mice most often focus on murine-relevant behaviors that are not necessarily applicable to the clinic. Similarly, mouse pathology is typically focused on invasive post-mortem analysis of tissues, processes that are not practiced in human patients. This lack of translatability renders many preclinical phenotypic characterizations difficult to translate to the clinic.

Comprehensive noninvasive biomarker panels are not available for many neurodegenerative disorders, and these are of special interest for pediatric disorders, as determining disease progression early on is critical to identifying proper clinical interventions and monitoring responses to potential corrective therapies. Batten disease (i.e., neuronal ceroid lipofuscinoses), a family of lysosomal storage disorders resulting from mutations in one of 13 genes, collectively represents the most common neurodegenerative disease in children (*2, 3*). Although the functions of many of these genes are unknown, much work in the past decade has been dedicated to developing and testing therapies. With the recent advancements in research tools, including high-throughput and high-content screening methods, the Batten disease scientific community has been progressing toward potential therapies at an unprecedented pace, and as a result, treatments are moving from the preclinical phase to clinical trials more quickly and efficiently than ever before. Also, with such heterogeneous disease states, resulting from different mutations that lead to more aggressive or protracted forms of the disease, clinical research teams are forming large, international collaborations to ensure that comprehensive natural history studies are completed and in place as a resource for all clinical trials. These interdisciplinary groups have paved the way for these natural history studies, led by the DEM-CHILD NCL Patient Database Consortium and the University of Rochester Batten Center (*4, 5*). Additionally, these groups have developed clinical rating scales to assess cognitive, motor, and behavioral function of patients with Batten disease (*6-8*). With a growing number of clinical trials for Batten disease therapies, there is an increasing need for noninvasive, clinically-relevant biomarkers to track therapeutic efficacy.

To address these needs, we used a mouse model of CLN6 Batten disease to perform an exhaustive multi-factorial characterization of biomarkers, moving away from mouse behavioral assays to clinical outcomes that are congruent to those used in human patients. CLN6 disease is caused by autosomal recessive mutations in *CLN6*, which results in the reduction or complete absence of the CLN6 protein. This disease is characterized by the accumulation of autofluorescent storage material in lysosomes, progressive neurodegeneration throughout cortex and thalamus, as well as massive gliosis throughout the central nervous system (CNS). Patients often present with language deficits, cognitive impairment and progressive motor decline. As the disease progresses, patients lose vision, develop seizures and ultimately succumb to the disease around 10-12 years of life. The spontaneously occurring *Cln6^nclf^* mouse model of CLN6 disease has been shown to faithfully recapitulate many of the hallmarks of the human disease both behaviorally and pathologically (*9, 10*), but noninvasive, clinically relevant assays have not yet been employed to characterize longitudinal changes in this model. Various scientists, our lab included, have performed a very comprehensive pathological assessment of Batten disease rodent and large animal models to reveal how different brain regions change over time, however, these experiments were all conducted on post-mortem brain samples (*11-20*). These studies have shown that brain pathology is present months before any noted behavioral changes, similar to what has been noted in human patients. Thus, cellular changes and degeneration are occurring in the the brain long before one notices any behavior or cognitive changes, so developing more sensitive tools for detecting disease states prior to the onset of behavioral symptoms would be of immense value in the clinic. Our previous work, as well as the work of others, has largely been focused on finding a single biomarker – the elusive “needle in a haystack” - associated with Batten disease and its progression (*21-24*), but this approach has failed to yield any reliable metrics.

In this study, rather than focusing on one or a few metrics, we used multiple imaging modalities as well as a comprehensive gait assessment to track hundreds of parameters over time, and performed a combinatorial analysis to identify a biomarker signature for CLN6-disease. Cohorts of wild type and *Cln6^nclf^* mice of both sexes were monitored longitudinally using noninvasive imaging, including T2-weighted magnetic resonance imaging (T2-MRI), diffusion tensor MRI (DTI), ^1^H magnetic resonance spectroscopy (MRS), and positron emission tomography (PET) was performed periodically between 3-12 months of age while kinematic gait analysis (KGA), which captures a large number of metrics describing gait, was assayed from 6-12 months of age. Variations of all of these techniques are widely used in the clinic and have shown correlations between mouse models and human subjects (*25, 26*). Once we had characterized these parameters in the CLN6 disease mouse model, we used a recently developed form of principal component analysis (PCA), contrastive PCA (cPCA), to cluster the four imaging modalities and gait analysis in order to derive new variables that best capture and define the progressive nature of the disease. Together, this approach provides a robust and translatable platform for longitudinal monitoring of disease progression that can have profound utility, not only for Batten disease, but in a variety of animal models of rare neuropediatric diseases.

## Results

### Progressive changes in brain volume and anatomy in a model of CLN6 disease

The selective vulnerability of various populations of neurons is a key feature of many neurodegenerative diseases, including Batten disease (*27, 28*). Prior studies have demonstrated that thinning of select anatomical regions and different cortical layers is present in *Cln6^nclf^* mice (*9, 12, 14, 19*), however, all of these measurements are based on invasive, histopathological analysis of post-mortem tissues. To identify progressive changes in brain architecture, we examined cohorts of mice longitudinally up to one year of age with T2 weighted magnetic resonance (T2-MRI) and diffusion tensor imaging (DTI).

Whole brain volume steadily decreased over time in *Cln6^nclf^* mice, with lower volume noted at 6, 9 and 12 months of age with clear differences in unique brain regions (Fig. 1). Progressive cortical atrophy began at 9 months and was profound at 12 months, where cortical volume was reduced in size by ~14%. Milder atrophy was also observed in the cerebellum at 12 months, culminating in a ~9% reduction in volume. Interestingly, there was a slight but significant increase in whole brain volume at 3 months of age (~1%). Historically, scientists have focused on one sex or mixed sexes for behavioral and pathological changes rather than analyzing them individually. Here we separated and tracked individual sexes, providing an innovative way of monitoring these animals over time and strengthening the translational utility. Sex differences were also evident, with the increase in whole brain volume at 3 months seen solely in male *Cln6^nclf^* mice, but decreasing in both sexes beginning at 6 months of age (**Fig. S1, Table S1**). Cortical atrophy also appeared earlier (at 9 months) in females than in males (at 12 months). In human patients and mouse models of neurodegenerative diseases, such profound cortical atrophy is typically accompanied by an increase in lateral ventricle volume (*29*). Surprisingly, we did not detect such changes in the *Cln6^nclf^* mice. Still, these results suggest progressive neurodegeneration results in multiple changes in brain structure volumes over the time course of the disease.

**Fig. 1.**
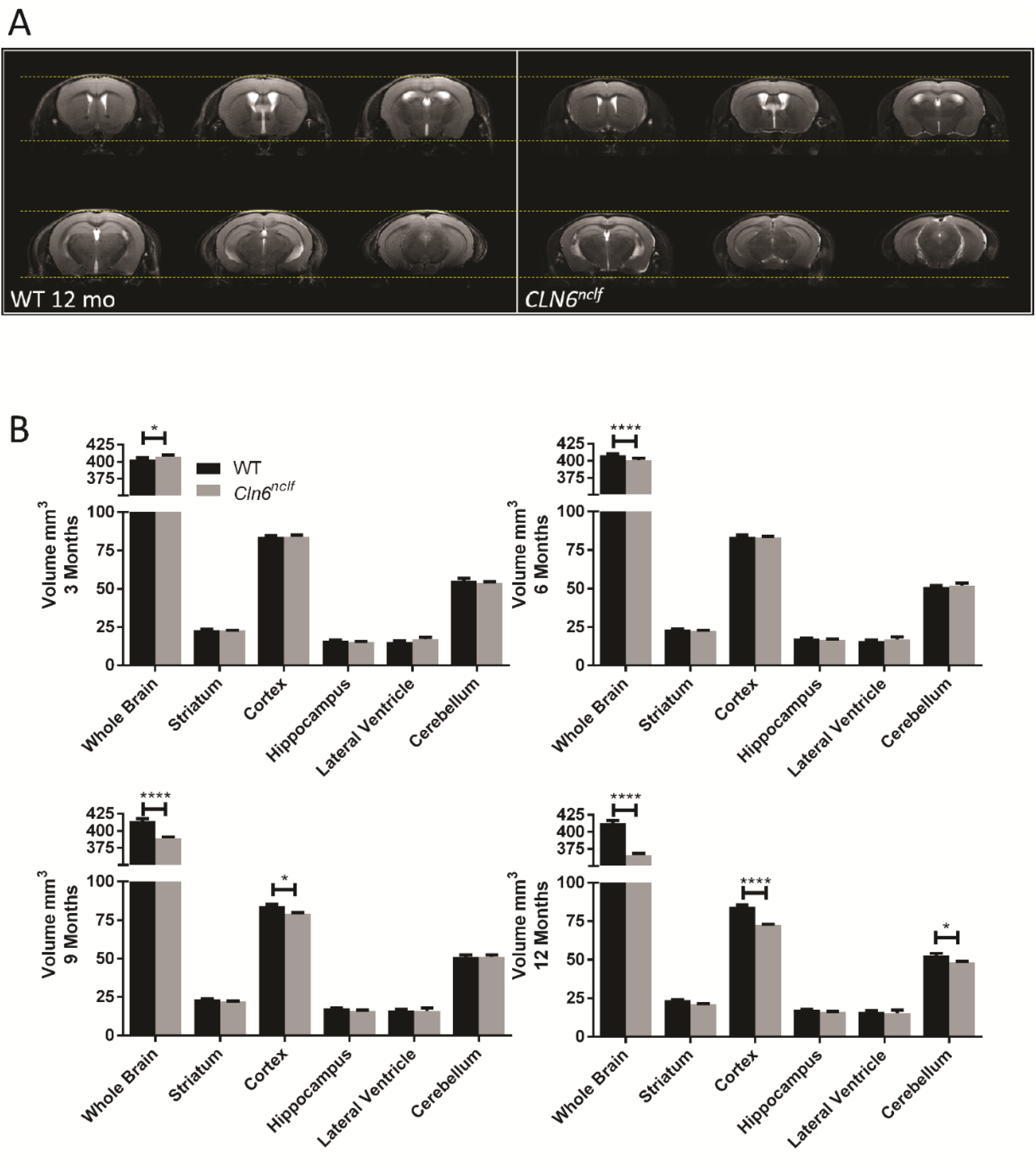
12 month old *Cln6^nclf^* mice have reduced brain volume in several key regions, as evidenced by MRI volumetry. A) Representative T2 weighted image panels for WT and *Cln6^nclf^* mice at the age of 12 months. Dashed yellow lines are to guide the eye for evident brain atrophy between the genotypes. B) Longitudinal brain structural changes over 3-12 month observation period. Progressive volume decline is seen in whole brain from 6 to 12 months, cortex volume at 9 and 12 months and cerebellum at 12 months. Data is mean ± SEM, n = 8 for WT (4 male, 4 female), n = 8 for *Cln6^nclf^* (4 male, 4 female). Statistical significances: unpaired, two-way ANOVA with Fisher’s LSD test. *p < 0.05, ****p < 0.0001.

To determine whether reductions in grey matter volume were accompanied by corresponding changes in white matter perturbations, we performed DTI to measure fractional anisotropy (FA) of several major CNS axon tracts. FA as measured by DTI reflects the level of preferred directionality of the diffusion of water molecules. Thus, higher FA values can reflect greater numbers, packing, or diameters of axons, lower variability in axon orientation, or more dense myelination (*30*). There were varying changes in FA of the forceps minor of the corpus callosum (fmi) over the time points monitored, and a consistent decrease in FA in the anterior portion of the anterior commissure (aca) beginning at 6 months of age (Fig. 2). Additionally, there were several regions with a decrease in FA at 12 months of age, correlating with the progressive and profound atrophy occurring at this time point. Within individual sexes, the FA of the fmi was increased in a male specific manner at 6 and 9 months of age (**Fig. S2**). Female *Cln6^nclf^* mice showed a decreased in FA in the splenium of the corpus callosum (scc) at 6 months of age, prior to their male counterparts, while both sexes began showing a decrease in FA in the aca beginning at 6 months of age (**Fig. S2, Table S2**). By 12 months of age, both sexes showed decreased FA in several white matter areas. These results demonstrate that progressive white matter defects are also prevalent in *Cln6^nclf^* mice, with unique sex specific perturbations in various white matter regions.

**Fig. 2.**
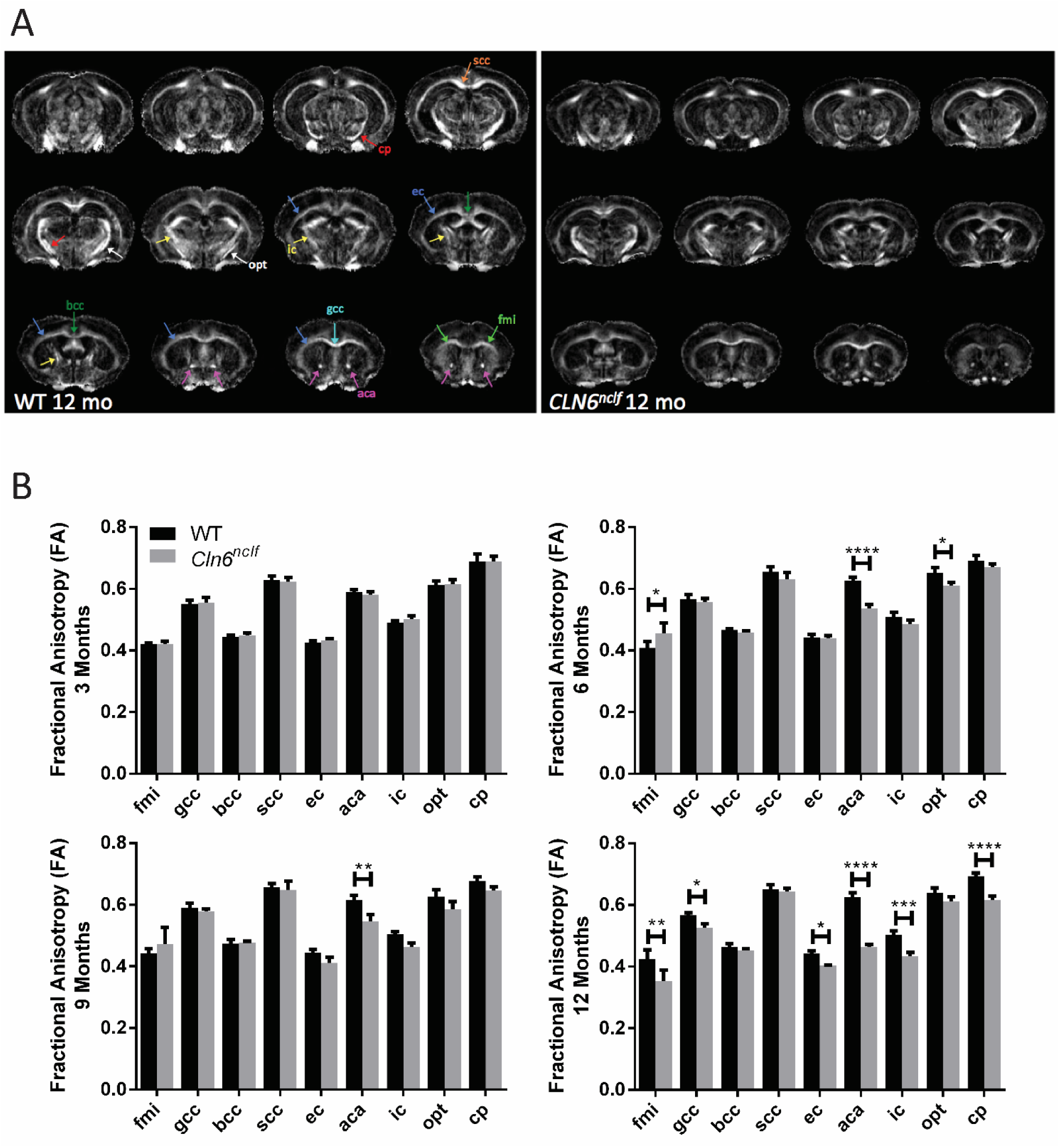
Diffusion tensor imaging identifies disrupted white matter architecture beginning at 6 months in *Cln6^nclf^* mice. A) Representative DTI image (fractional anisotropy) panels for WT and *Cln6^nclf^* mice at the age of 12 months. B) Longitudinal fractional anisotropy changes over 3-12 month observation period. Progressive white matter declines were detected in the aca beginning at 6 months of age in *Cln6^nclf^* mice, while various white matter regions showed reduced volumes at 12 months of age. Data is mean ± SEM, n = 8 for WT (4 male, 4 female), n = 8 for *Cln6^nclf^* (4 male, 4 female). Statistical significances: unpaired, two-way ANOVA with Fisher’s LSD test. *p < 0.05, **p < 0.01, ***p < 0.001, ****p < 0.0001. Abbreviations: fmi – forceps minor of corpus callosum, gcc – genu of corpus callosum, bcc – body of corpus callosum, scc – splenium of corpus callosum, ec – external capsule, aca – anterior commissure (anterior part), ic – internal capsule, opt – optic tract, cp – cerebral peduncle.

### Alterations in brain metabolism, metabolites, and markers of brain inflammation associated with CLN6 disease

Abnormalities in brain metabolism in the form of decreases in glucose uptake and increases in various markers of neuroinflammation is a common signature of a number of neurodegenerative disorders (*31, 32*). To explore whether such changes are present in the *CLN6-*Batten disease mouse model, we performed ^1^H magnetic resonance spectroscopy (MRS) and positron emission tomography (PET) imaging. MRS was first used to examine longitudinal changes in various brain metabolites in the frontal cortex. While we observed subtle changes in a variety of metabolites at various time points, the most significant changes were observed for glutamine (GLN), N-acetylaspartate (NAA), and NAA + N-acetylaspartylglutamic acid (NAA+NAAG) (Table 1). GLN levels steadily increased in *Cln6^nclf^* mice over time, reaching significantly elevated levels at 12 months of age in both sexes (Table 1). NAA and NAA+NAAG levels steadily decreased over time, reaching significantly lower levels at 9 and 12 months of age in both sexes. Additionally, taurine (TAU) was reduced at 3 – 9 months, more prominently in male mice, and creatine (Cr) + phospho-creatine (PCr) reduced at 3 months in female mice, though these differences resolved over time. These results suggest that alterations in glutamate-glutamine cycling and excitatory signaling may be a prominent feature of this disease model, mirroring what has been found in other studies(*33*). Furthermore, NAA decreases we observed mirror those seen in human patients with a number of neurodegenerative disorders including Batten disease (*34-36*), suggesting that this may be a useful marker for monitoring neurodegeneration in *Cln6^nclf^* mice.

**Table 1.**
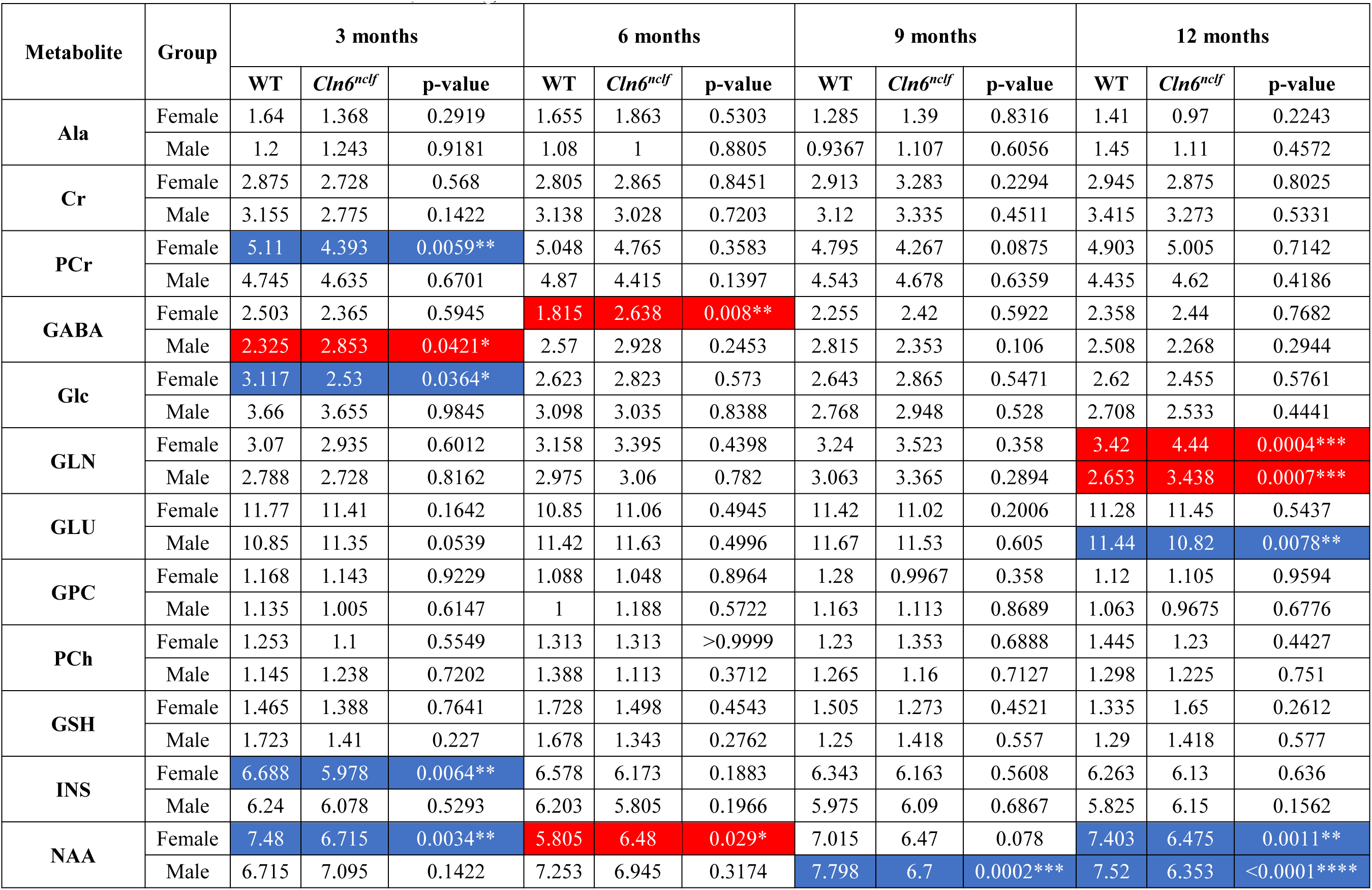
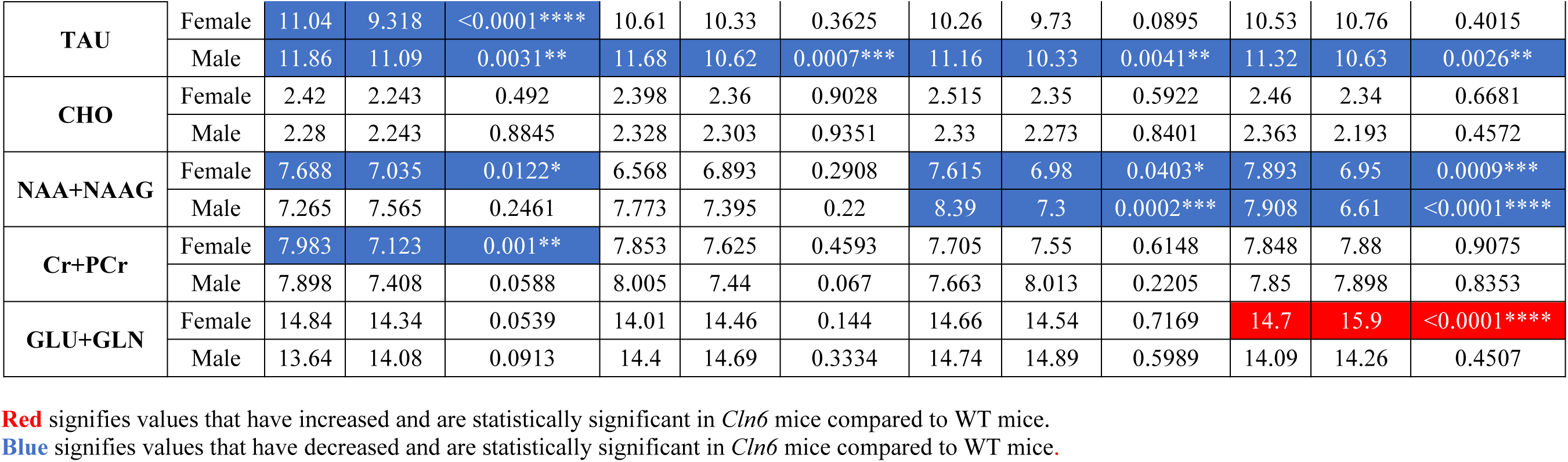
^1^H-MRS: Metabolite Concentration (mmol/kg)

To explore potential changes in brain metabolism (i.e. brain glucose utilization), we used PET to monitor uptake of fluorodeoxyglucose (^18^F, FDG), a PET-detectable proxy for glucose. At 12 months, where we observed the most severe alterations in brain anatomy, we found that FDG standardized uptake values (SUV) were significantly compromised in all regions examined in male *Cln6^nclf^* mice and in many regions in female *Cln6^nclf^* mice (Table 2). We also utilized PET to measure the uptake of ^18^F-FEPPA. This ligand binds the translocator protein (TSPO), which is upregulated in microglia in response to neuroinflammation, one of the earliest reported pathological changes in Batten disease mouse models (*37*). Interestingly, at 13 months of age, uptake was reduced in male mice in the basal forebrain and septum (BFS), cortex, and olfactory bulb while increased uptake was found in females in the central gray matter, superior colliculi, and thalamus (**Table S3**). It must be noted, however, that we observed significantly decreased body weights in male and female *Cln6^nclf^* mice at this time point (**Fig. S3**). Since SUV is negatively correlated with body weight, this complicates any comparison of SUV values between groups. These results, obtained at the advanced stage of 13 months of age, suggest that finding significant differences in ^18^F-FEPPA uptake at earlier time points may be even more challenging. Taken together, the differences we observed could be indicative of alterations in glucose uptake, brain perfusion, neuronal metabolism, and inflammatory status.

**Table 2.**
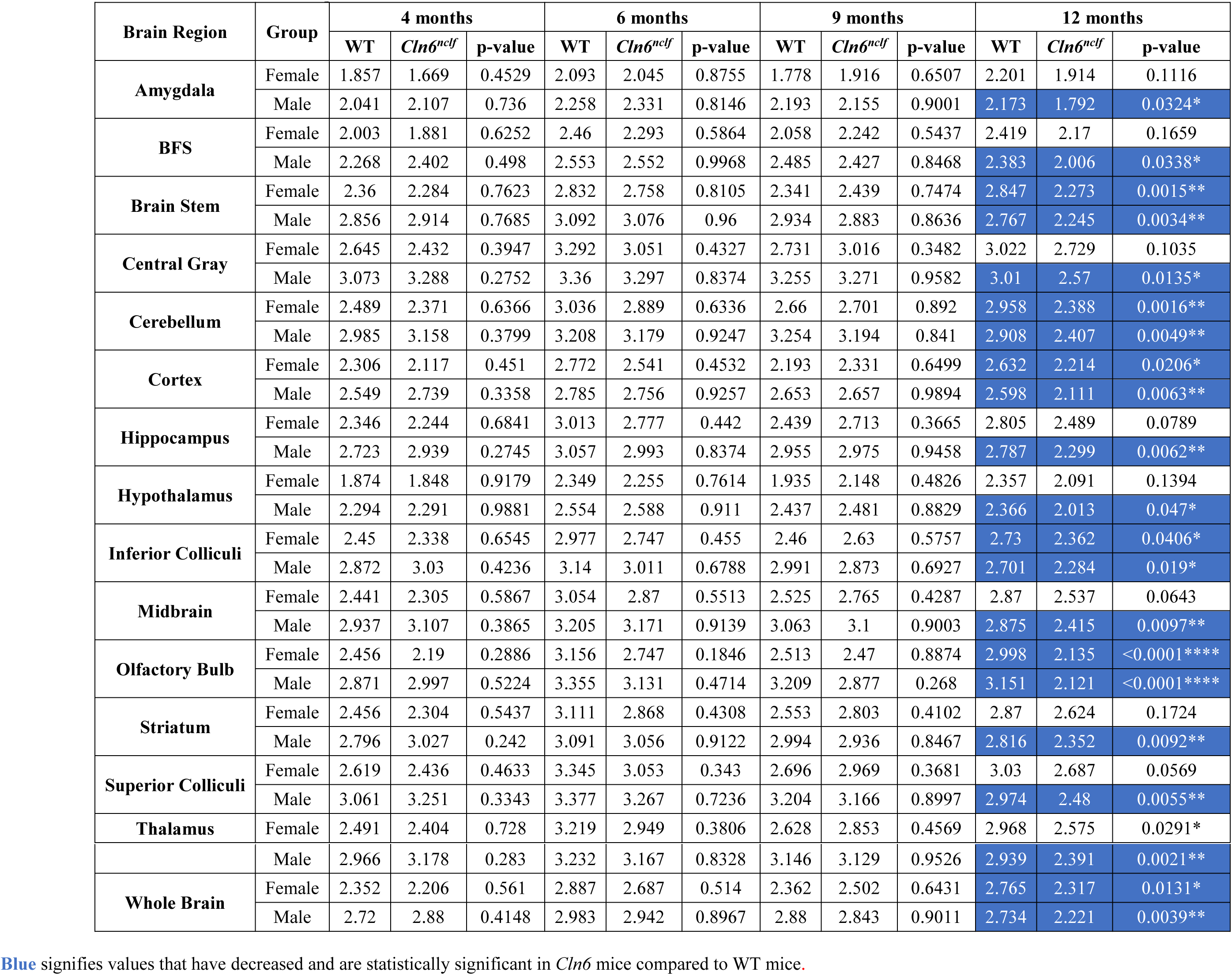
FDG-PET: ^18^F-FDG Standard Uptake Values

### Kinematic Gait Analysis identifies novel motor disturbances in Cln6^nclf^ mice

The profound anatomical and metabolic abnormalities we observed in *Cln6^nclf^* mice would be expected to lead to behavioral disturbances. While motor coordination deficits have been identified in *Cln6^nclf^* mice using crude measures such as rotarod performance (*9, 38*), an exhaustive analysis of gait parameters that have good human correlates in the clinic has not been conducted. We used kinematic gait analysis (KGA) to examine 97 gait parameters longitudinally in the *Cln6* mutant mice (**Table S4, Fig. S4-S9**). This included a detailed analysis of gait cycle, body and head orientation and positioning, and multiple fore and hind limb parameters. Additionally, we used a PCA based approach to calculate overall gait scores.

Many individual parameters varied between genotypes and time points, but several predominant gait features defined the *Cln6^nclf^* gait. *Cln6^nclf^* mice ambulated with slower overall speed, decreased durations of diagonal gait mode (trotting), increased durations of double support, slower overall speed, lower hind body posture reflected in lower tail base, hip and iliac crest heights and decreased knee and ankle heights, and elevated tow clearance in both fore and hind limbs (**Fig. S4**). Different gait features, which are manifested in sets of highly correlating parameters, can be identified using principal component analysis [PCA; (*39*)]. PCA is a commonly used technique to reduce the dimensionality of multivariate data sets (*40, 41*), or more specifically, to determine a few linear combinations of the original variables that can be used to summarize the data set while retaining as much information as possible. Moreover, the use of PCA also enables the inspection and identification of the mutual correlations between the original kinematic parameters. PCA based calculation of overall gait scores revealed profound and progressive changes in *Cln6^nclf^* mice (Fig. 3). The overall score is a weighted average of the gait variables using weights from the discriminant vectors, which emphasizes which kinematic parameters are contributing to the overall score (Fig. 3B). When analyzing pooled sexes, *Cln6^nclf^* gait scores were significantly different from controls at all time points, culminating in an approximately 12-fold increase at 12 months of age. When analyzing sexes individually, individual sex score in males increased significantly beginning at 6 months of age. In females, the differences was significant only at 12 months of age. Additionally, there was a significant genotype by age interaction in gait scores, supporting differences in gait over time across the two genotypes (Fig. 3A). These results demonstrate that *Cln6* mutant mice have profound, progressive gait disturbances, mirroring phenotypes that have been described in human CLN6 disease patients (*42, 43*).

**Fig. 3.**
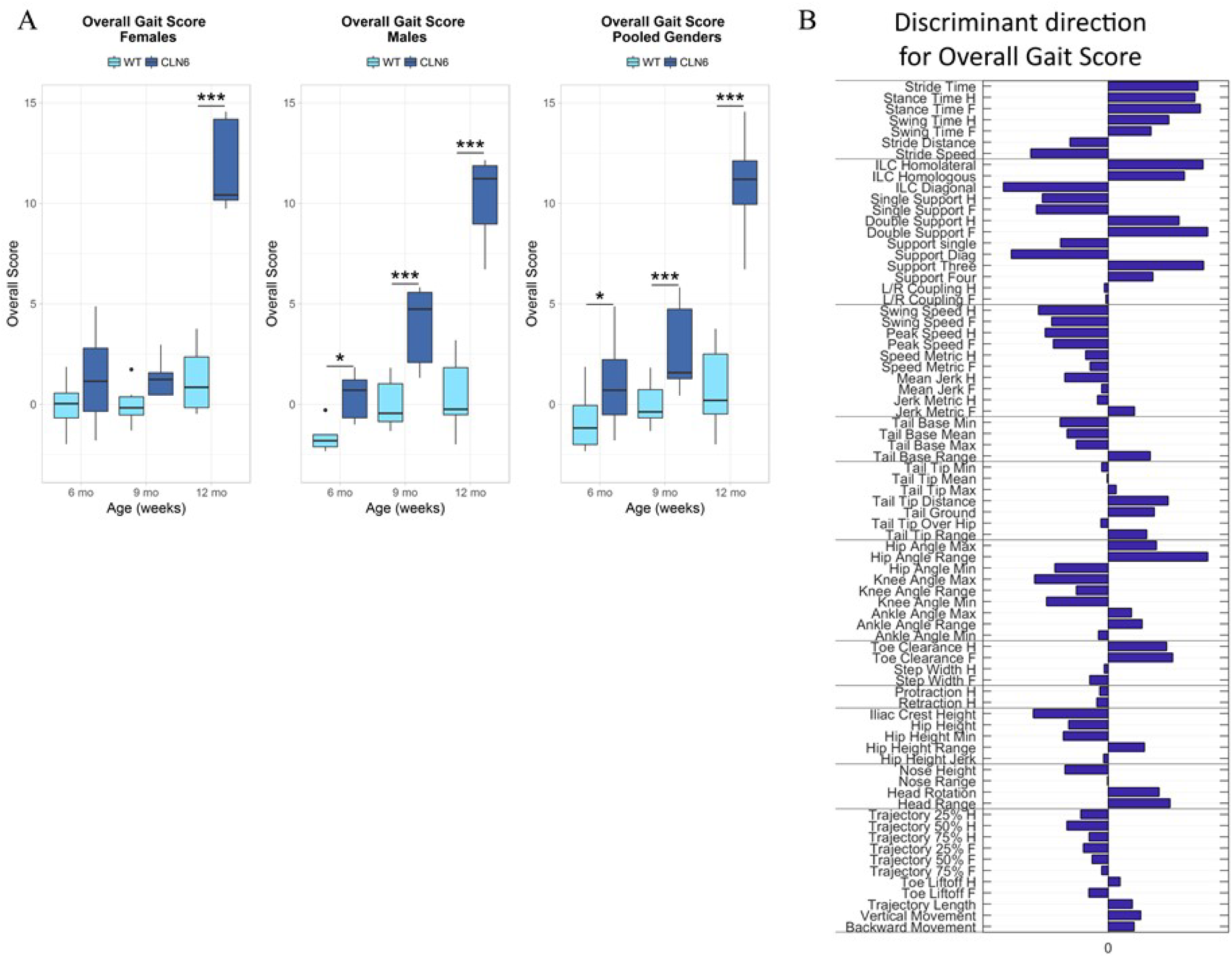
Kinematic gait analysis followed by PCA identifies progressive alterations in gait. A) The overall score is based on the principal component score differences between the *Cln6^nclf^* and the WT groups in 63 selected kinematic parameters altogether. The score can be interpreted as “how far away is an individual from the average WT towards the direction of the average *CLN6^nclf^*?”. The mean of the WT group score is equal to zero. B) The discriminant direction bar graph illustrates how each kinematic parameter is weighted in the score. The bar graph also represents an overall kinematic fingerprint of the *Cln6^nclf^* model over the three time points: zero level correspond to average WT. Data is presented as group means +/-SEM. n = 12 for WT (6 male, 6 female), n = 11 for *CLN6^nclf^* (5 male, 6 female). Statistical significances: unpaired t-test \**p* < 0.05, \*\**p* < 0.01, \*\*\**p* < 0.001, \*\*\*\**p* < 0.0001

### Contrastive PCA identifies the core progressive symptomatology in Cln6^nclf^ mice

Where other studies or natural history studies have fallen short is that they try to look at each individual parameter in isolation rather than looking at the system as a whole. We performed contrastive PCA (cPCA) in an effort to holistically capture the progressive changes present in *Cln6^nclf^* mice. Our phenotypic analysis identified a number of novel anatomical, metabolic, and behavioral phenotypes associated to the *Cln6* mutant animals. This large volume of data presented an important question: *Is there a core set of phenotypes that strongly correlate with one another and best describe the progressive nature of the Cln6^nclf^ disease course?* Identifying such a set of phenotypes would be of great utility in utilizing this model for not only preclinical testing of novel therapies but could have profound clinical value. So instead of looking for ‘a single needle in a haystack,’ we analyzed ‘the whole haystack’ using statistical methods capable of reducing the data without losing any information.

To perform this analysis, we used a recently developed statistical technique, contrastive PCA (cPCA) (*44*). This procedure identifies a low dimensional structure that is enriched in one genotype over another, thus calculating cPC scores that accentuate differences between genotypes. Our analysis included MRI volumetry from 6 brain regions, DTI from 9 white matter regions, MRS for 19 metabolites in the PFC, FDG-PET from 15 brain regions, and KGA for 97 parameters, longitudinally from both genotypes of mice. First, we calculated 1st level principal components (PCs), three for each modality. This reduced the original 144 variables yielding 15 unique variables (PCs), many of which highlighted progressive changes in *Cln6^nclf^* mice (Fig. 4).

**Fig. 4:**
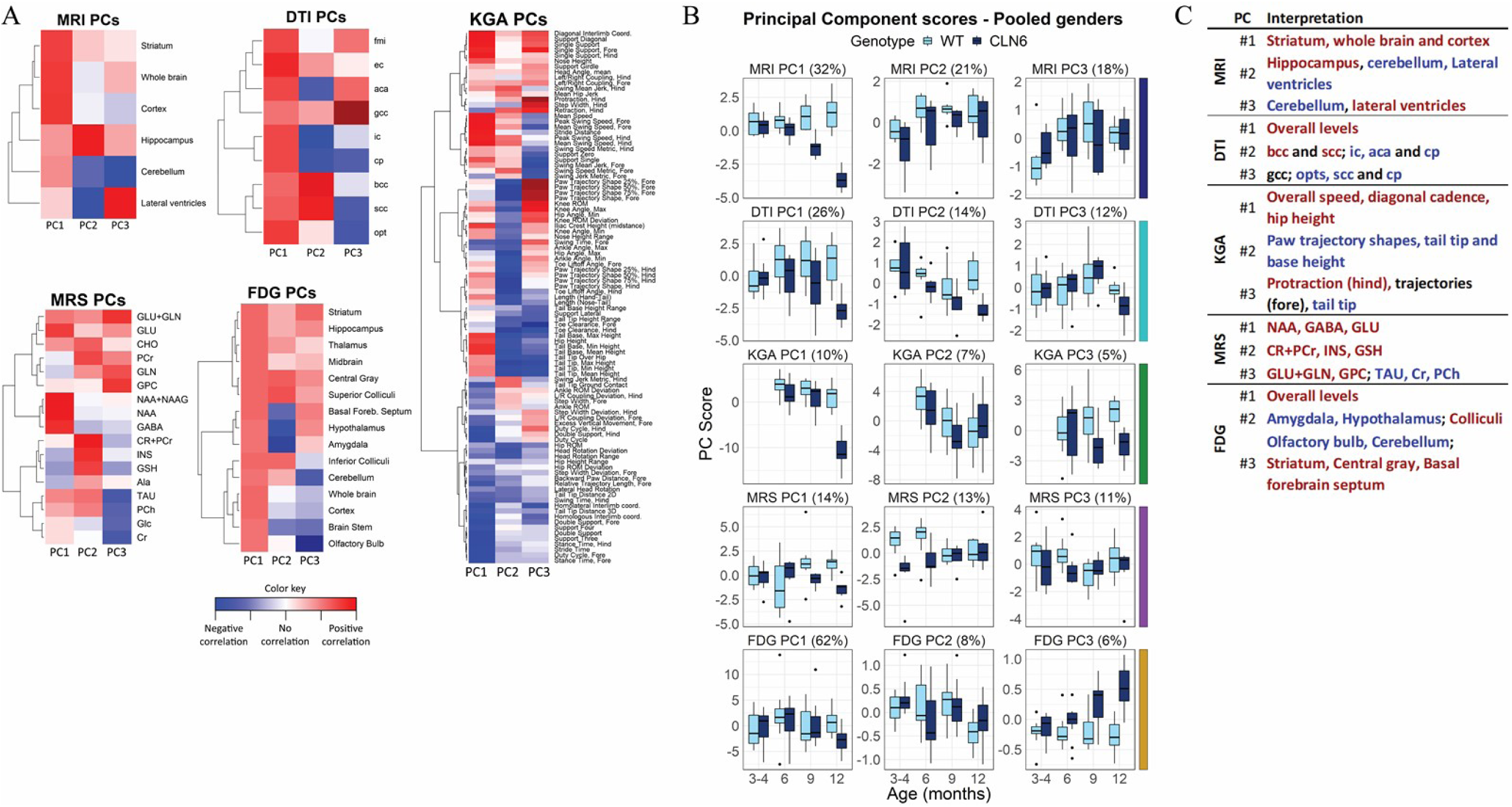
Principal component analysis demonstrates how the most influential combinations of non-invasive imaging and gait analysis variables contribute to the progressive changes in the *Cln6^nclf^* disease state. A) Three first principal components of each five modalities, shown as heatmaps, illustrate the identified correlation structure within each dataset. Strong cell color, red or blue, indicates that the variable has high loading in the corresponding PC. Moreover, a set of variables with high loadings within the same PC are strongly correlated. The correlation between variables of same color is positive (red or blue), and negative in case of opposite colors (red and blue). For example, in MRI PC1, volumes of striatum, whole brain and cortex are correlated directly. In MRI PC3, cerebellum and lateral ventricles correlate inversely. B) Interpretations of the principal components are based on findings which variables are most strongly correlated within each component. C) Phase 1 principal component scores. Data is presented as boxplots. The percentage numbers indicate variance explained by the corresponding PC.

Next, we used cPCA to calculate 2nd level cPC scores based on the 1st level PCs (Fig. 5). The top cPC score, cPC1, included contributions primarily from MRI, FDG-PET, DTI, and KGA, demonstrating that gait performance was associated with anatomical and metabolic changes in various brain regions (Fig. 5A). cPC1 proved to be very effective at characterizing phenotypic progression in *Cln6^nclf^* mice (Fig. 5B). The top 5 features (i.e., the five 1st level PCs with highest contribution) in the cPC1 were: 1) MRI PC1 (overall brain volume), 2) KGA PC1 (overall speed + diagonal cadence + overall hip height), 3) FDG PC3 (increase in striatum, central grey matter, and basal forebrain associated with decrease in cerebellum and olfactory bulb.), 4) DTI PC1 (overall level), and 5) DTI PC2 (increase in the body of the corpus callosum and splenium of the corpus callosum, decrease in anterior commissure, internal capsule and cerebral peduncle). Taken together, the decreased cPC1 score in *Cln6* mutant mice can be interpreted as decrease in structural brain volume (MRI PC1) associated with decrease in overall mobility (KGA PC1), and metabolic changes (FDG PC3), and reductions in overall (DTI PC1) and local (DTI PC2) anisotrophy. Remarkably, the genotype difference was significant at each of the time points examined for pooled sexes, as well as each sex separately. The second cPC, cPC2, highlighted a much wider variability in *Cln6^nclf^* mice at all time points. The variation in cPC2 is mostly explained by difference between sexes within *Cln6* mutant animals. There was a significant difference between *Cln6^nclf^* male versus *Cln6^nclf^* female at all time points. The variation in cPC2 is mostly explained with MRI PC2 (hippocampus (↑), cerebellum/lateral ventricles (↓)) and MRI PC3 (cerebellum (↓), lateral ventricles (↑)) (Fig. 5A). However, partially due to this variability, differences between genotypes were mostly insignificant (Fig. 5B). Still, cPC1 provides a powerful summary of the *Cln6^nclf^* phenotype starting early in disease, and could be the basis for highly sensitive assays for future therapeutic testing. Thus, the data from this comprehensive analysis could be used for generating a final disease score, given the longitudinal differences between genotypes. Such a score could in turn be used to determine efficacy of potential treatments in clinical trials.

**Fig. 5:**
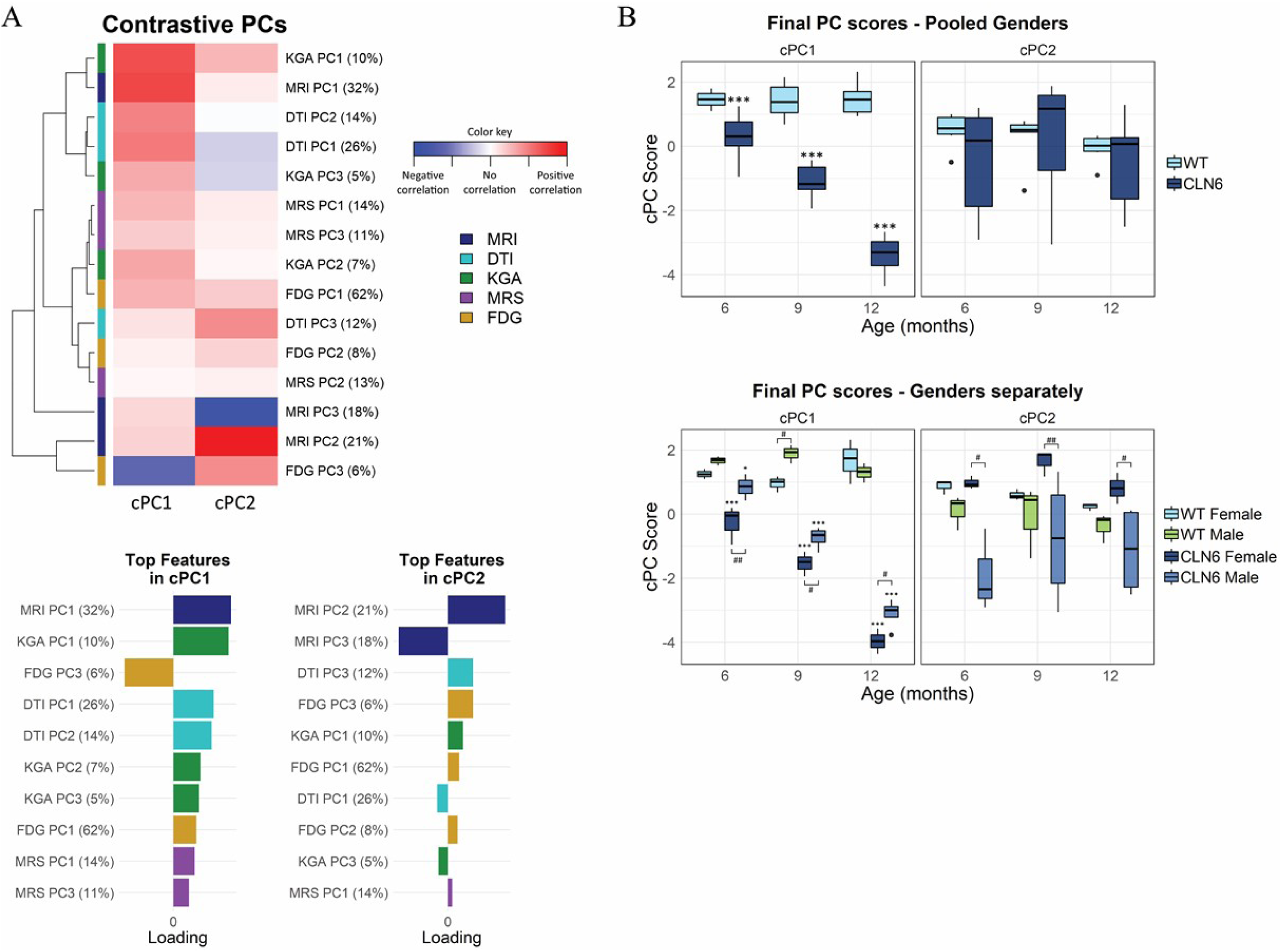
Contrastive principal component analysis defines new variables that best capture the progressive changes present in the *Cln6^nclf^* disease state. A) Contrastive principal components shown as heatmap. The most strongly correlated phase 1 components in the cPCs are presented as color coded bar graphs. B) Final principal component scores presented as boxplots, pooled sexes (top) and sexes separately (bottom). The first component, most emphasized by MRI PC1, KGA PC1, FDG PC1 and DTI PC1 and 2, demonstrate progressively increasing phenotype difference (*) in both sexes. The cPC2, consisting mostly of MRI PC1 and 2 but also DTI PC3 and FDG PC3, reveals sex difference (#)in *Cln6* mice at all ages. n = 6 for WT (3 male, 3 female), n = 7 for C*ln6^nclf^* (4 male, 3 female). Statistical significances: unpaired t-test of estimated mean differences *p < 0.05, **p < 0.01, ***p < 0.001

## Discussion

Pediatric neurodegenerative disorders are typically diagnosed by a combination of clinical assessment, neuroradiologic imaging, cellular pathology, and genetic testing (*45-47*). Unfortunately, even with advancements in genomic diagnostics, seldom are children definitively diagnosed following initial clinical assessment, and more typically, numerous misdiagnoses are offered before a correct diagnosis (*48*). The clinical features of Batten disease, are a combination of cognitive dysfunction, dementia, retinopathy, and seizures (*49-52*) but can vary depending on the subtype of the disease and the precise genetic mutation the parent has. Common phenotypic features include muscular hypotonia, microcephaly, myoclonus, epilepsy, ataxia, behavioral changes, visual decline, and brain atrophy (*29, 51-55*). Although there have been reported sex differences in Batten disease patients, scientists have historically focused on one sex or mixed sexes for behavioral and pathological changes. By separating and tracking individual sexes, this study provides an innovative way of monitoring these animals over time, strengthening the translational utility. Our findings reinforce that the *Cln6^nclf^* model recapitulates many aspects of the human disease course, including the progressive nature, of CLN6 disease. We show that *Cln6^nclf^* mice have profound and progressive changes in brain anatomy and metabolism, and that these changes correlate strongly with abnormalities in gait parameters. cPC1, which encapsulates the changes that appeared most consistently and with the greatest magnitude, shows very large and increasing differences between wild type and *Cln6^nclf^* mice from 6-12 months of age. In addition to providing insights into the pathological manifestations of *Cln6* deficiency, this work suggests a number of clinical assessments that may be most useful in assessing CLN6 patients. Although our genetically identical mice differ in many important ways from a more heterogeneous population of human patients, the beauty of this system is that it isn’t reliant on one metric to produce the score of disease state, and can still detect progressive changes in the presence of between-individual variance.

In Batten disease patients, general brain structure changes include cerebral and cerebellar atrophy, callosal thinning, enlarged ventricular space, white matter maturation, and thalamic density, which may appear abnormal before any clinical symptoms appear (*29, 53, 55-63*). White matter and basal ganglia of the thalami display significantly decreased signal intensity T2-weighted MR images in addition to the increased signal of the periventricular matter (*56, 58, 60, 64, 65*). Single photon emission computed tomography studies show hypoperfusion in cerebral and cerebellar cortices, most prominent in anterior frontal, posterior temporoparietal and occipital cortices, and abnormal lesions can be detected before structural abnormalities appear (*66-69*). MRS has shown reductions in N-acetyl aspartate (NAA), glutamine + glutamate + GABA (Glx), creatine (Cr) and choline (Cho) compounds, and elevation of lactate, lipids, and myoinositol (mIns) (*53, 60, 61, 70, 71*). Additionally, PET imaging show dysfunction of nigrostriatal dopaminergic neurons (reduced [^18^F]fluorodopa uptake) (*72*), reduced metabolism beginning in the calcarine area followed by a widespread decrease throughout the cortex and thalamus (decreased glucose utilization in FDG-PET) (*73*), and impaired striatal neuronal function (reduced striatal dopamine D1) (*74*).

The functional imaging modalities employed here revealed significant changes in the diseased mice that also correlate with similar changes reported for Batten disease patients. The *Cln6^nclf^* mice display generalized whole brain atrophy starting at 6 months of age that rapidly progresses, and a reduction in brain metabolites and metabolism as measured by MRI, MRS and FDG-PET, respectively. Region-specific volumetry changes included decreased volume of the cortex and cerebellum. Additionally, there was an opposite trend between male and female diseased mice for lateral ventricle volume, with males showing an increase in ventricular volume at 3 months that was stabilized through 12 months of age, while females showed a steady decrease in ventricular volume over time. These results may be in part due the general decrease in brain volume for both sexes. MRS identified a severe reduction in NAA at 9 – 12 months for both males and females. FDG-PET revealed highly reduced glucose metabolism in all brain regions at 12 months of age. Diffusion tensor MRI and fractional anisotropy showed that brain changes in *Cln6^nclf^* mice involve not only grey matter, but white matter as well. Contrary to many neurodegenerative diseases where corpus callosum changes are often pronounced, deeper white matter structures such as anterior commissure, external and internal capsule seem to be primarily affected in *CLN6^nclf^* mice although corpus callosum changes become evident with disease progression. Decrease of FA is normally attributed to e.g. demyelination, axonal loss/disruption or incoherence due to pathological processes. Unfortunately, supporting NCL related histology-verified DTI research, both clinical and pre-clinical, is still largely lacking. Our data takes important steps in providing observations from animal models of NCL that can be utilized to justify the use of DTI more often in a neuroradiological assessment of Batten disease.

Although motor coordination deficits have been identified in *Cln6^nclf^* mice using behavioral tests such as rotarod performance (*9, 38*), traditionally, there have been significant inter-and intra-lab variability in animal behavior testing. Variation in behavior results have depended on genetic background of the mouse model, genetic drift, breeding strategies, geographical location of testing facility, variability in behavior protocols implemented, gender of the animal handler, animal diet, and water quality (*75, 76*). As for differences in behavior testing protocols, different labs implement various training time and number of repeated measures. Fine kinematic gait analysis conducted on 97 unique measurements identified unique movement scores among the *Cln6^nclf^* mice. The overall gait score is based on differences between the wild type and *Cln6^nclf^* groups in all the PC scores. Ultimately, the overall kinematic effects of a pharmacological agent may be seen in a highly sensitive manner with a wide therapeutic window, which suggests that this tool will be of great value for further preclinical and translational studies in various neurodegenerative disease.

Our results demonstrate the translational power of an exhaustive, multi-factorial characterization of biomarkers in the *Cln6^nclf^* mouse model of CLN6 disease. Our characterization suggests which clinical tools may be most useful in monitoring patients, and also provides a highly sensitive platform for testing therapies in mice. We focused on two main categories of tools which have direct correlates in the clinic: kinematic gait analysis (KGA) and noninvasive neuroimaging. KGA monitors ambulating mice, capturing a large number of metrics that together describe deviations from normal gait. This method is widely employed in the clinic to objectively describe gait abnormalities in patients. While there are large differences in ambulation mode between bipedal humans and quadrupedal mice, recent developments have demonstrated correlations between parameters of human and mouse gait, and the translational utility of gait analysis in mouse models (*25*). Similarly, in terms of noninvasive brain imaging, there are many anatomical similarities between the human and murine brain, and studies have shown that noninvasive imaging often detects similar changes in human patients and mouse models of neurodegenerative disease (*26*).

Furthermore, many researchers have focused on single blood-, saliva-, and cerebrospinal fluid-based biomarkers in mice and large animal models of Batten disease (*21-23, 77-80*). These focused studies have led us to conclude that no singular target will provide a reliable biomarker of Batten disease. Additionally, one particularly important study analyzed autopsy brains and cerebrospinal fluid (CSF) from deceased Batten disease patients and identified numerous significant changes of potential biomarkers of disease (*81*). However, this study was conducted on post-mortem brain samples and CSF, which would represent samples that are either impossible to analyze in living patients or invasively acquired. We sought to go beyond simply identifying a single biomarker or group of phenotypes that are present in this mouse model, and to provide a more useful characterization that also identifies which phenotypes correlate most closely with one another, and which phenotypes best demonstrate the progressive nature of the disease. To accomplish this goal, we used cPCA, which is highly effective for identifying the parameters that best define the differences between groups. This analysis is intended to identify cPCs that amplify the differences between healthy and disease subjects, thus increasing the sensitivity to detect changes, positive or negative, in disease progression. Such a metric could have great power for identifying potentially useful therapies in preclinical studies in mice, large animal models, or for quantifying progression or therapeutic benefit in patients, and removes the barrier of looking for a single biomarker of disease by combining multiple biomarkers into one clinical score of disease state/progression.

The strength and novelty of this study stems from utilizing multiple non-invasive functional imaging modalities to longitudinally track and compare wild type and diseased animals over time, and combining the noninvasive imaging, including T2-MRI, DT, MRS, and PET, with KGA in a cPCA to generate a disease score. This score identifies diseased animals and provides a robust and translatable platform for long term monitoring of animal models of disease that can also be applied to large animal models of neurodevelopmental and neurodegenerative diseases as well as clinical patients. The platform we have developed will be widely applicable to the study of a variety of animal models of neurodegenerative disease. In addition to identifying novel phenotypes of these disorders, this combination of neuroimaging, behavior, and statistical analysis should enable the identification of cPCA-based phenotypes that accentuate progressive changes, greatly enhancing the power of these models for preclinical studies. Furthermore, the translational nature of the techniques this platform utilizes may provide important insights for clinicians regarding which noninvasive imaging and behavioral modalities may be most useful in the diagnosis and ongoing assessment of patients with neurodegenerative diseases. Establishing these parameters for patients with neurodegenerative diseases is imperative for future drug screening and utility in human clinical trials. For CLN6 disease patients currently enrolled in the phase I/II clinical trial (ClinicalTrials.gov Identifier: NCT02725580) (*82*), there is currently no way to conclusively monitor responses to these treatments over time. Having these capabilities in place would allow for quantitative analysis of patient responses. Additionally, in the future these metrics may be combined with existing data from blood-and CSF-based biomarkers to develop a comprehensive panel of noninvasive biomarkers for many neurodegenerative diseases.

## Materials and Methods

### Study Design

Neuroimaging, gait analysis, and principal component analyses were conducted on aged-matched wildtype and *Cln6* mutant mice (description of animals in Ethics Statement/Animals) to determine a longitudinal biomarker scoring system in a preclinical model of neuropediatric disease. We hypothesized that by looking at various non-invasive disease markers as a system, rather than individually, we could provide a highly sensitive tool that may be translatable to the clinic. Sample size, endpoints, and rules for stopping data collection were determined based on our previously published studies on this model (*83*). No outliers were removed from any data sets. All animal experiments were performed as specified in the license authorized by the national Animal Experiment Board of Finland and according to the National Institutes of Health (Bethesda, MD, USA) guidelines for the care and use of laboratory animals. Experiments were conducted in an AAALAC accredited laboratory. Animals’ care was in accordance with institutional guidelines. 3-13 month, male and female wild type (WT) and homozygous *Cln6*-mutant mice (*Cln6^nclf^*; JAX stock #003605) on C57BL/6J backgrounds were utilized for all studies, were housed under identical conditions, and all experimenters were blinded to genotype.

### Magnetic Resonance Imaging and Spectroscopy

MRI experiments were performed using a horizontal 11.7T magnet with a bore size of 160 mm, equipped with a gradient set capable of maximum gradient strength of 750 mT/m and interfaced to a Bruker Avance III console (Bruker Biospin GmbH, Ettlingen, Germany). A volume coil (Bruker Biospin GmbH, Ettlingen, Germany) was used for transmission and a surface phased array coil for receiving (Rapid Biomedical GmbH, Rimpar, Germany). Mice were anesthetized using isoflurane, fixed to a head holder and positioned in the magnet bore in a standard orientation relative to gradient coils. Temperature of the animals were monitored and maintained between 36-37°C throughout the experiments. To avoid prolonged anesthesia/study day, MR experiments were performed in two separate scanning sessions; 1) MRI volumetry and localized 1H-MR spectroscopy from frontal cortex (total duration appr. 45 minutes) and 2) diffusion tensor imaging (DTI, total duration appr. 1 hour).

Structural MRI was performed with a standard Turbo-RARE sequence with TE_eff_ of 34 ms (RARE factor of 8), TR of 3150 ms and 8 averages. Thirty-one 0.45 mm slices were collected with field-of-view of 20×20 mm^2^ and 256×256 matrix (78 microns in-plane resolution). Region of interest analysis was performed in MATLAB (Mathworks Inc., Natick, MA, USA) environment observer blinded for study groups. Whole brain, cortex, striatum, hippocampus, lateral ventricle and cerebellar volumes were analyzed.

For the acquisition of proton MRS data, frontal cortex voxel (2.2×1.6×1.8 mm3, 6.3 μl localized volume) was selected based on structural MR images described above. Automatic 3D gradient echo shimming algorithm was used to adjust B0 homogeneity in the voxel. The water signal was suppressed using variable power RF pulses with optimized relaxation delays (VAPOR) to obtain B1 and T1 insensitivity. A PRESS sequence (TE = 10 ms) combined with outer volume suppression (OVS) was used for the pre-localization. Data were collected by averaging 512 excitations (frequency corrected for each FID) with TR of 2 s, number of points 2048 and spectral width of 5 kHz. Excitation frequency was shifted −2 ppm, to minimize the chemical shift phenomenon within the selected voxel. In addition, a reference spectrum without water suppression (NT=8) was collected from the identical voxel using the same acquisition parameters. Peak areas for resolved metabolites were analyzed using LCModel (Stephen Provencher Inc., Oakville, Canada) using >CRLB 20% as exclusion criterion for individual metabolites within analyzed spectrum.

Diffusion tensor MRI (DTI) was performed using 4-segment EPI sequence with 30 diffusion directions (TE/TR =23.5/4000 ms, b-values 0 and 970 s/mm2). Field-of-view of 12.80 × 10.24 mm2 (with saturation slice) was used with matrix of 160 × 128, resulting 80 microns in-plane resolution. Fifteen 0.6 mm slices were acquired with 6 averages. Preprocessing of DTI-data consisted eddy-current correction and brain masking. Diffusion tensor was calculated using FSL (https://fsl.fmrib.ox.ac.uk/fsl/) and resulting fractional anisotropy maps were processed with manual ROI-analysis in the MATLAB environment (Mathworks Inc.) for the following anatomical structures; forceps minor of the corpus callosum (fmi), genu of corpus callosum (gcc); body of corpus callosum (bcc); splenium of corpus callosum (scc); external capsule (ec); anterior commissure anterior part (aca); internal capsule (ic); optic tract (opt); cerebral peduncle (cp).

### Longitudinal ^18^F-FDG and ^18^F-FEPPA PET Imaging

The C*ln6^nclf^* and wild type mice were longitudinally PET scanned at the age of 4, 6, 9 and 12 months. After an overnight fasting, to standardize blood glucose levels, the mice were injected intravenously with a 150 µl bolus of ^18^F-FDG in sterile saline, 14.0 ± 1.5 MBq. The mice were anesthetized with isoflurane 20 minutes after the injection and positioned into small animal PET/CT (BioPET/CT, BioScan, USA). Three mice were scanned simultaneously in list mode for 25 minutes, 30 minutes post the ^18^F-FDG injection. PET scan was followed by CT scan for anatomical orientation and CT-based attenuation map for image reconstruction. The sinograms were reconstructed with 3D-OSEM, 1 iteration and 25 subsets, with attenuation correction. Image analysis was performed with PMOD software (PMOD Technologies, Switzerland, v.3.7). The PET images were co-registered with mouse brain MRI template and standardized uptake values (SUVs) were calculated on different brain regions.

At the age of 13 months the *Cln6^nclf^* and wild type mice were anesthetized with isoflurane, cannulated into lateral tail vein and positioned into small animal PET/CT (BioPET/CT, BioScan, USA). Dynamic 90.5 minute PET scan was started and 150 µl of [^18^F] FEPPA, 11.4 ± 1.0 MBq, was injected intravenously 30 seconds after the start of the scan. PET scan was followed by CT scan for anatomical orientation and CT-based attenuation map for image reconstruction. The sinograms were reconstructed with 3D-OSEM, 1 iteration and 25 subsets, with attenuation correction. Image analysis was performed with PMOD software (PMOD Technologies, Switzerland, v.3.7). The PET images were co-registered with mouse brain MRI template and standardized uptake values (SUVs) were calculated on different brain regions from last 30 minutes of the PET scan.

### Fine motor kinematic analysis

Mice were subjected to kinematic gait analysis test at 6, 9, and 12 months of age, using a Motorater apparatus (TSE-Systems GmbH, Bad Homburg, Germany) designed for the assessment of fine motor skills in rodents. The equipment consists of a brightly illuminated Plexiglas corridor (153 × 5 × 10 cm) under which is situated a high-speed camera. Prior to the test, the mice were shaved under light isoflurane anesthesia, and the essential body points, such as joints and tail, were marked for tracking. The gait performance data was captured using a camera operated 300 frames per second, imaging the gait simultaneously from three different views (underside and both sides). The movement was analyzed from the three views, first using the Simi Motion software (Simi Reality Motion Systems GmbH, Unterschleissheim, Germany). Approximately 5-6 complete strides were analyzed from each mouse. Only strides with continuous ambulatory movement were included in the data. The raw kinematic data thus comprised of the movements of 24 different body points in coordinates related to the ground. Different gait patterns and movements were analyzed using a custom made automated analysis software, resulting 97 distinctive kinematic gait parameters(*84*) (**Fig. S7-S12**) such as: general gait pattern parameters (e.g., stride time and stride speed, step width, stance time and swing time during a stride, interlimb coordination, etc.), body posture and balance parameters (e.g., toe clearance, iliac crest height, hip height, hind limb pro-, and retraction, tail position, tail movements), and fine motor skills (e.g. swing speed during a stride, jerk metric during swing phase, angle ranges and deviations of distinct joints, vertical and horizontal head movements).

### Principal Component Analysis (PCA)

PCA is a linear transformation based on principal component coefficients and eigenvectors. The transformed, new, uncorrelated variables are called the principal components (PC). The first PC corresponds to such linear combination of data which has the largest possible variance. The second PC has again the largest possible variance of what is left when the proportion of the first PC is discarded, and so on for the rest of the PCs. The total number of PCs was selected using Kaiser’s rule, i.e., only those PCs were retained which explain more (variance) than one normalized original variable. The eigenvectors also reveal information about the internal structure of the data, i.e., mutually correlated parameters. Each PC score is emphasized by different combination of mutually correlated original variables. So, each PC is a linear combination of original variables, such that some variables are emphasized in one PC and some other variables in another. Variables which are emphasized in a PC are mutually correlated. On the other hand, the PCs themselves are totally uncorrelated.

The Overall Gait Score is based on differences between the wild type and the *Cln6^nclf^* groups in all the PC scores. Thus, the purpose of that score is to identify a combination of original variables, a “fingerprint”, which characterizes the disease model in the best possible way and differentiates the two groups. After the “fingerprint”, or discriminant direction vector, has been constructed, the overall gait analysis scores are obtained by projecting the (normalized) parameter data of each individual mouse onto the discriminant direction vector. The average wild type group individual has always score equal to zero, and roughly half of the controls have always negative score, reflecting that their overall gait performance is in the opposite direction than the performance of the disease model phenotype. Moreover, the average disease model phenotype has always a positive score value, and the magnitude of that value corresponds to magnitude of phenotype specific deviations in an overall gait pattern.

The cPCA is based on PCA of two datasets, X and Y, where X (target) may consist of data of different genotypes, ages, treatments and sexes, and Y (background) consists of control group data, not including variation due to genotype or disease model. The aim in cPCA is to create a subset combination of original parameters, which simultaneously 1) maximizes variance in X (interesting *and “*universal” features) and 2) minimizes variance in Y (“universal” features only). In other words, the cPCA is used to discover the combination of those interesting variables, which all together add contrast between the background and the target, i.e., genotype specific variables.

### Statistical Analysis

Individual tests were run in Graphpad Prism (Ver 7.04). In general, an ordinary two-way ANOVA was used at each time point, using genotype and brain region/metabolite as main factors. An uncorrected Fisher’s LSD test was used to determine statistical significance in individual brain regions/metabolites across genotypes. Specific statistical tests and sample size are described in the figure legends. Raw data and complete data tables with exact p-values are available from the authors upon reasonable request. Graphs as presented as Mean +/-SEM, *p<0.05, **p<0.01, ***p<0.001, ****p<0.0001. PCA was performed and cPCA was implemented and performed in R according to Abid et al. (*44*): A language and environment for statistical computing (Version 3.5.0, R Foundation for Statistical Computing, Vienna, Austria).

## Supplementary Materials

Figure S1: Longitudinal brain structural changes in sex separated data

Figure S2: Longitudinal fractional anisotropy changes in sex separated data

Figure S3: Body weight analysis at 13 months

Figure S4: Spatiotemporal kinematic parameters

Figure S5: Interlimb coordination parameters

Figure S6: Kinematic parameters describing body posture, toe clearance, hind limb protraction and retraction, nose height and head rotation

Figure S7: Kinematic parameters describing limb trajectory profiles and excess movements during swing phase of gait

Figure S8: Kinematic parameters describing tail tip movements

Figure S9: Hip, knee and ankle angles

Table S1: MRI volumetry values

Table S2: Diffusion tensor imaging: fractional anisotropy values

Table S3: FEPPA-PET: ^18^F-FEPPA standard uptake values

Table S4: Kinematic gait parameters and definitions

## Acknowledgments: Funding

This work was supported by funding to JMW NIH R01NS082283 as well as institutional support from Sanford Research and Charles Rivers..

## Author contributions

K.K.L, A.N and J.M.W conceptualized and designed the studies. D.T. bred and shipped the mice. K.K.L J.R., and T.B. conducted the experiments and generated data. T.B.J., J.J.B., K.K.L, J.T.C, K.A.W., T.B., J.R. T.B., A.N., and J.M.W. critically analyzed and plotted data. T.B.J and J.J.B wrote the manuscript. T.B.J, J.J.B, K.K.L, J.T.C, K.AW, T.B, J.R., T.H., M.V., J.T.P., A.N, and J.M.W. reviewed and edited the manscript.

## Data and materials availability

All data generated in this study can be found in the paper or supplemental materials or provided upon request.

