## Supplemental Information for "A multimodal approach to identify clinically relevant parameters to monitor disease progression in a preclinical model of neuropediatric disease"

**Supplementary Materials:**

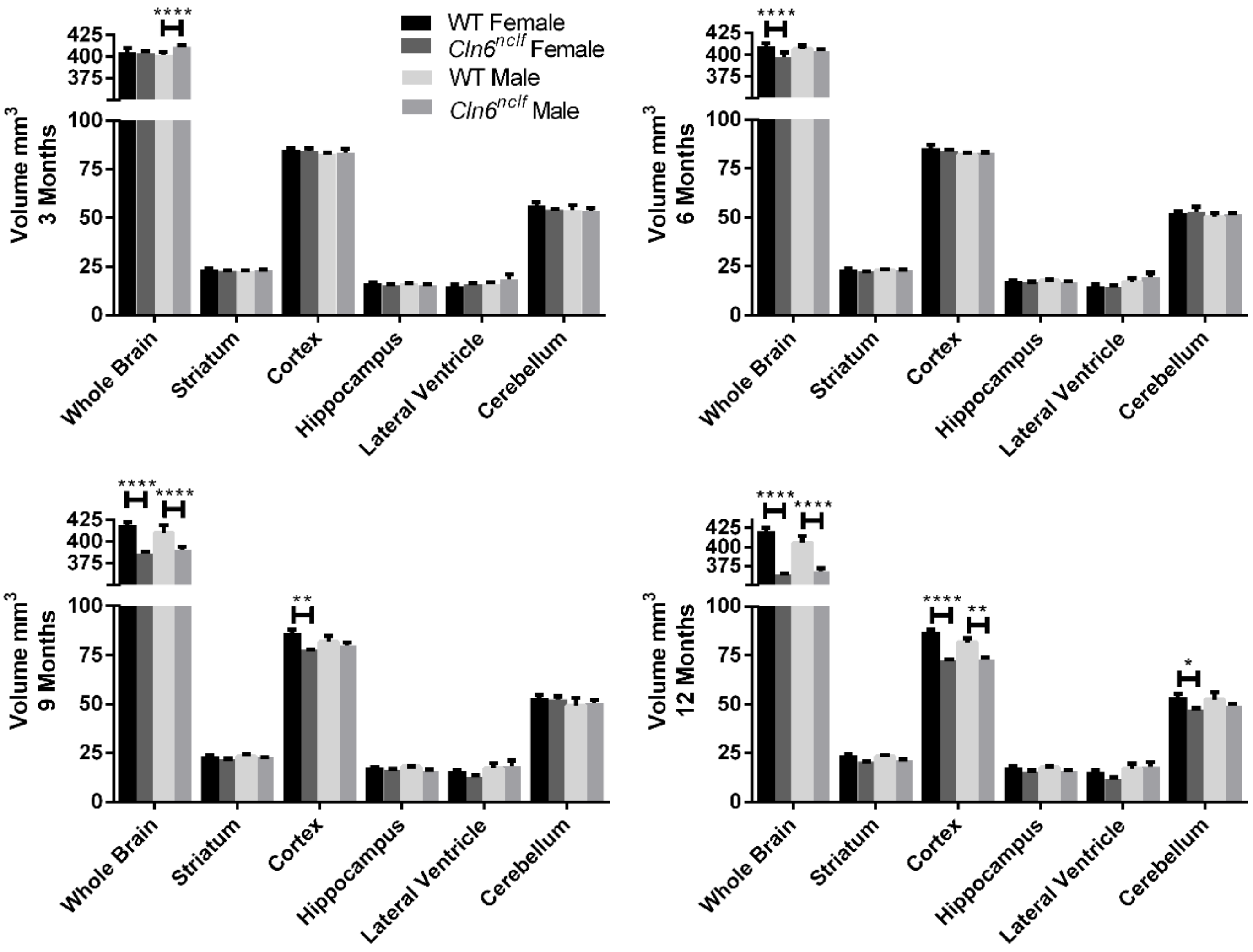

**Fig. S1: Longitudinal brain structural changes in sex separated data.** Brain structural changes over 3-12 months observation period in sex separated data for *Cln6*$^{nclf}$. Data is mean ± SEM, n = 8 for WT (4 male, 4 female; data pooled), n = 4 for *Cln6*$^{nclf}$ for both male and female mice. Statistical significances: unpaired, two-way ANOVA with Fisher's LSD test. P-values can be found in Table S1.

**Table S1: MRI volumetry (mm$^3$) values**

| Brain Region | Group | 3 months | | | 6 months | | | 9 months | | | 12 months | | |
|---|---|---|---|---|---|---|---|---|---|---|---|---|---|
| | | WT | $Cln6^{nclf}$ | p-value | WT | $Cln6^{nclf}$ | p-value | WT | $Cln6^{nclf}$ | p-value | WT | $Cln6^{nclf}$ | p-value |
| **Whole Brain** | Female | 405.1 | 404.1 | 0.6818 | 409.9 | 397.6 | <0.0001**** | 418.9 | 386.4 | <0.0001**** | 420 | 363.5 | <0.0001**** |
| | Male | 401.3 | 411.7 | <0.0001**** | 406.9 | 404.5 | 0.3236 | 409.8 | 390.7 | <0.0001**** | 405.3 | 368.3 | <0.0001**** |
| **Striatum** | Female | 23.76 | 22.72 | 0.6796 | 23.64 | 22.26 | 0.5652 | 23.26 | 21.73 | 0.5887 | 23.91 | 20.56 | 0.2458 |
| | Male | 22.52 | 22.92 | 0.8752 | 23 | 23 | >0.9999 | 23.35 | 22.63 | 0.7812 | 23.22 | 21.59 | 0.5379 |
| **Cortex** | Female | 84.9 | 84.41 | 0.8422 | 85.32 | 83.76 | 0.5172 | 86.41 | 77.66 | 0.0028** | 87.05 | 72.09 | <0.0001**** |
| | Male | 83.17 | 83.65 | 0.8469 | 81.85 | 82.73 | 0.7146 | 81.69 | 80.01 | 0.523 | 81.55 | 72.85 | 0.0016** |
| **Hippocampus** | Female | 16.46 | 15.55 | 0.7159 | 17.28 | 16.86 | 0.8606 | 17.6 | 16.36 | 0.6617 | 17.58 | 15.6 | 0.4907 |
| | Male | 16.36 | 15.32 | 0.6775 | 17.76 | 16.65 | 0.6425 | 17.92 | 15.87 | 0.4347 | 17.56 | 16.05 | 0.57 |
| **Lateral Ventricle** | Female | 15.04 | 15.77 | 0.7695 | 15.09 | 14.33 | 0.7537 | 15.76 | 12.66 | 0.276 | 15.52 | 11.71 | 0.1865 |
| | Male | 15.76 | 18.63 | 0.2533 | 16.73 | 19.48 | 0.2547 | 17.12 | 18.59 | 0.573 | 16.99 | 18.04 | 0.6926 |
| **Cerebellum** | Female | 56.72 | 54.35 | 0.3444 | 52.21 | 52.51 | 0.9016 | 52.86 | 52.24 | 0.8258 | 53.56 | 47.34 | 0.0331* |
| | Male | 54.11 | 53.63 | 0.8461 | 50.09 | 51.67 | 0.5126 | 49.13 | 50.48 | 0.6065 | 52.28 | 48.96 | 0.2138 |

**Red** signifies values that have increased and are statistically significant in *Cln6* mice compared to WT mice.

**Blue** signifies values that have decreased and are statistically significant in *Cln6* mice compared to WT mice.

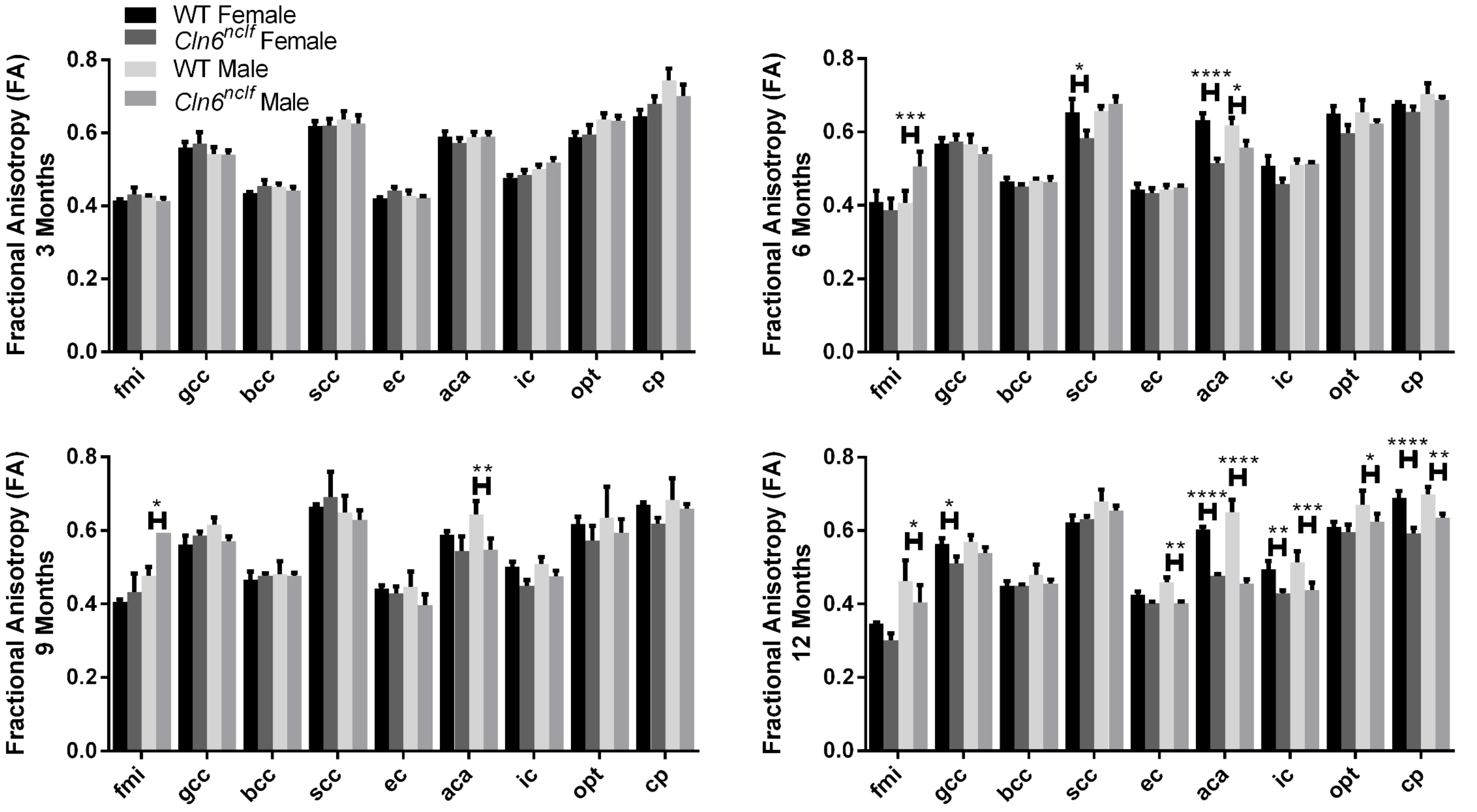

**Fig. S2: Longitudinal fractional anisotropy changes in sex separated data**. Fractional anisotropy changes over 3-12 months observation period in sex separated data for *Cln6*$^{nclf}$. Data is mean ± SEM, n = 8 for WT (4 male, 4 female; data pooled), n = 4 for *Cln6*$^{nclf}$ for both male and female mice. Statistical significances: unpaired, two-way ANOVA with Fisher's LSD test. P-values can be found in Table S2.

**Table S2: Diffusion tensor imaging: fractional anisotropy values**

| Brain Region | Group | 3 months | | | 6 months | | | 9 months | | | 12 months | | |
|---|---|---|---|---|---|---|---|---|---|---|---|---|---|
| | | **WT** | ***Cln6^nclf*** | **p-value** | **WT** | ***Cln6^nclf*** | **p-value** | **WT** | ***Cln6^nclf*** | **p-value** | **WT** | ***Cln6^nclf*** | **p-value** |
| **Fmi** | Female | 0.4146 | 0.4319 | 0.5069 | 0.4095 | 0.386 | 0.4405 | 0.4058 | 0.4318 | 0.4386 | 0.3464 | 0.3001 | 0.1306 |
| | Male | 0.4277 | 0.4139 | 0.5712 | 0.407 | 0.5063 | 0.0006*** | 0.4772 | 0.5945 | 0.0185* | 0.4632 | 0.4043 | 0.0273* |
| **Gcc** | Female | 0.5602 | 0.5702 | 0.6599 | 0.5685 | 0.5745 | 0.8313 | 0.5625 | 0.5856 | 0.492 | 0.5629 | 0.5104 | 0.0258* |
| | Male | 0.5421 | 0.5407 | 0.9491 | 0.5663 | 0.5398 | 0.3478 | 0.6167 | 0.5714 | 0.147 | 0.5693 | 0.5383 | 0.1518 |
| **Bcc** | Female | 0.4355 | 0.4552 | 0.3828 | 0.466 | 0.4515 | 0.6069 | 0.4657 | 0.4769 | 0.7382 | 0.4499 | 0.4487 | 0.9593 |
| | Male | 0.4514 | 0.4415 | 0.6611 | 0.4668 | 0.4643 | 0.9293 | 0.4812 | 0.4772 | 0.8983 | 0.4797 | 0.4559 | 0.2703 |
| **Scc** | Female | 0.619 | 0.6195 | 0.9822 | 0.6533 | 0.5838 | 0.0239* | 0.6653 | 0.691 | 0.4992 | 0.6228 | 0.6308 | 0.7324 |
| | Male | 0.6376 | 0.6259 | 0.6043 | 0.6568 | 0.677 | 0.4727 | 0.6481 | 0.6281 | 0.5187 | 0.6787 | 0.6542 | 0.2566 |
| **Ec** | Female | 0.4211 | 0.4427 | 0.3397 | 0.443 | 0.433 | 0.7226 | 0.4423 | 0.4295 | 0.7031 | 0.4243 | 0.4026 | 0.3517 |
| | Male | 0.4282 | 0.4218 | 0.7741 | 0.443 | 0.448 | 0.8591 | 0.447 | 0.3974 | 0.1118 | 0.4599 | 0.4032 | 0.0097** |
| **Aca** | Female | 0.59 | 0.5725 | 0.4388 | 0.6323 | 0.5153 | <0.0001**** | 0.5881 | 0.5434 | 0.1843 | 0.603 | 0.4762 | <0.0001**** |
| | Male | 0.5882 | 0.5896 | 0.9508 | 0.6185 | 0.5568 | 0.0302* | 0.6428 | 0.5479 | 0.0028** | 0.6501 | 0.455 | <0.0001**** |
| **Ic** | Female | 0.4772 | 0.4849 | 0.7308 | 0.5075 | 0.4575 | 0.078 | 0.5011 | 0.4497 | 0.1271 | 0.4934 | 0.4282 | 0.006** |
| | Male | 0.5015 | 0.5192 | 0.4316 | 0.5113 | 0.5135 | 0.9363 | 0.5091 | 0.4755 | 0.2803 | 0.513 | 0.438 | 0.0007*** |
| **Opt** | Female | 0.5874 | 0.5948 | 0.7437 | 0.6505 | 0.5963 | 0.0562 | 0.618 | 0.5725 | 0.1767 | 0.6097 | 0.5953 | 0.5361 |
| | Male | 0.6378 | 0.6333 | 0.8424 | 0.6538 | 0.6225 | 0.2686 | 0.6351 | 0.5946 | 0.1937 | 0.67 | 0.6242 | 0.0353* |
| **Cp** | Female | 0.6459 | 0.6797 | 0.1351 | 0.6763 | 0.6548 | 0.4786 | 0.6695 | 0.6191 | 0.1867 | 0.6888 | 0.5915 | <0.0001**** |
| | Male | 0.7446 | 0.7003 | 0.0902 | 0.704 | 0.6873 | 0.5524 | 0.6833 | 0.6594 | 0.4418 | 0.6982 | 0.6346 | 0.0039** |

**Red** signifies values that have increased and are statistically significant in *Cln6* mice compared to WT mice.
**Blue** signifies values that have decreased and are statistically significant in *Cln6* mice compared to WT mice.

**Table S3. FEPPA-PET: $^{18}$F-FEPPA standard uptake values**

| Brain Region | Group | WT | *Cln6*$^{nclf}$ | p-value |
|---|---|---|---|---|
| **Amygdala** | Female | 0.6899 | 0.6984 | 0.8433 |
| | Male | 0.9012 | 0.8007 | 0.1041 |
| **BFS** | Female | 0.5427 | 0.5458 | 0.9422 |
| | Male | 0.7708 | 0.6073 | 0.0085** |
| **Brain Stem** | Female | 0.81 | 0.8897 | 0.0658 |
| | Male | 0.9868 | 0.9823 | 0.9428 |
| **Central Gray** | Female | 0.491 | 0.5845 | 0.031* |
| | Male | 0.6771 | 0.6296 | 0.4423 |
| **Cerebellum** | Female | 0.7268 | 0.7404 | 0.7529 |
| | Male | 0.9284 | 0.8631 | 0.2904 |
| **Cortex** | Female | 0.526 | 0.5435 | 0.6852 |
| | Male | 0.7258 | 0.5956 | 0.0356* |
| **Hippocampus** | Female | 0.5232 | 0.5861 | 0.1462 |
| | Male | 0.7005 | 0.634 | 0.2815 |
| **Hypothalamus** | Female | 0.6571 | 0.6561 | 0.9818 |
| | Male | 0.8137 | 0.7062 | 0.0826 |
| **Inferior Colliculi** | Female | 0.5617 | 0.616 | 0.2087 |
| | Male | 0.7352 | 0.6716 | 0.3031 |
| **Midbrain** | Female | 0.559 | 0.6296 | 0.1031 |
| | Male | 0.7394 | 0.6708 | 0.267 |
| **Olfactory Bulb** | Female | 0.6534 | 0.6459 | 0.8612 |
| | Male | 0.9062 | 0.7107 | 0.0017** |
| **Striatum** | Female | 0.4952 | 0.5531 | 0.1811 |
| | Male | 0.7038 | 0.6038 | 0.1058 |
| **Superior Colliculi** | Female | 0.4778 | 0.5656 | 0.0428* |
| | Male | 0.6704 | 0.6069 | 0.3038 |
| **Thalamus** | Female | 0.4855 | 0.5783 | 0.0323* |
| | Male | 0.6557 | 0.6189 | 0.551 |
| **Whole Brain** | Female | 0.6022 | 0.6392 | 0.392 |
| | Male | 0.798 | 0.7068 | 0.1402 |

**Red** signifies values that have increased and are statistically significant in *Cln6* mice compared to WT mice.
**Blue** signifies values that have decreased and are statistically significant in *Cln6* mice compared to WT mice

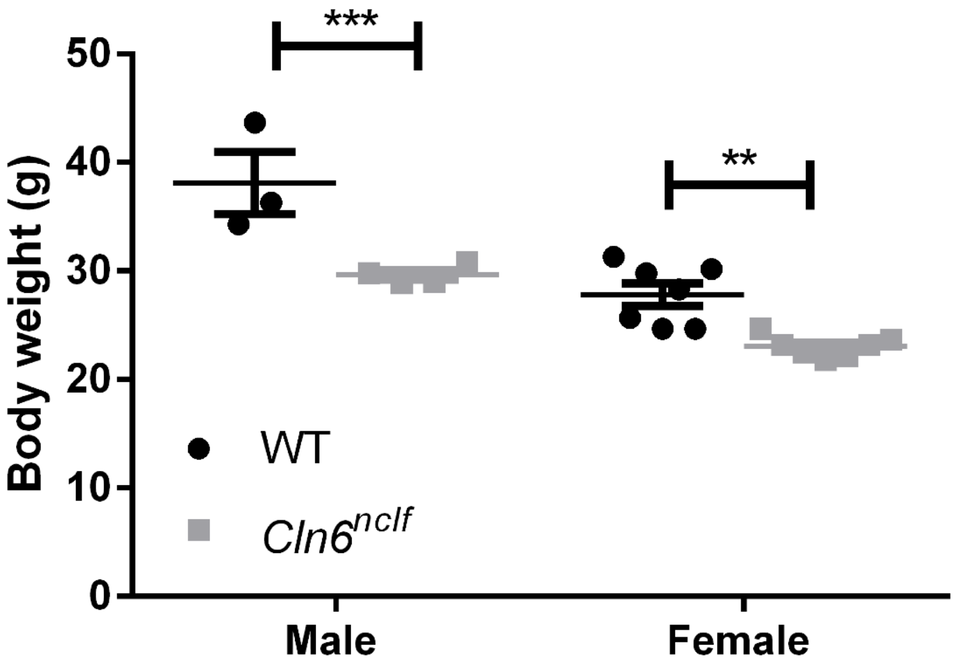

**Fig. S3: Body weight analysis at 13 months**. Differences in body weight complicates SUV value calculations from $^{18}$F-FEPPA images. Data is mean ± SEM, n = 10 for WT (3 male, 7 female), n = 11 for *Cln6$^{nclf}$* (4 male, 7 female). Statistical significances: unpaired, two-way ANOVA with Fisher's LSD test.

**Table S4: Kinematic gait parameters and definitions**

| | Parameter | Definition |
|---|---|---|
| | **Stride Time** | Duration of a full stride. |
| **Spatio-temporal** | **Mean Speed** | Mean ambulatory movement speed. |
| | **Stride Distance** | Distance moved during a full stride. |
| | **Stance Time (hind, fore)** | Duration the paw is in contact with the floor, stance phase. |
| | **Swing Time (hind, fore)** | Duration the paw is in the air, swing phase. |
| | **Mean Swing Speed (hind, fore)** | Mean paw movement speed during swing phase. |
| | **Peak Swing Speed (hind, fore)** | Maximum paw movement speed during swing phase. |
| | **Swing Speed Metric (hind, fore)** | Ratio of the Mean Swing Speed to Peak Swing speed. |
| | **Mean Swing Jerk (hind, fore)** | The degree of non-smoothness, i.e., rate of acceleration change, of a paw during middle half of swing phase. |
| | **Swing Jerk Metric (hind, fore)** | Ratio of the Swing Mean Jerk to Swing Peak Speed. A normalized swing trajectory smoothness parameter. |
| | **Duty cycle (hind, fore)** | Percentage of stride time the limb is in contact with the floor. |
| | **Homolateral Interlimb Coordination** | Proportion of whole stride duration in which ipsilateral paws are simultaneously in stance or swing phase. |
| **Interlimb Coordination** | **Homologous Interlimb Coordination** | Proportion of whole stride duration ipsi- and contralateral fore or hind paws are simultaneously in stance or swing phase. Pace. |
| | **Diagonal Interlimb Coordination** | Proportion of whole stride distance in which a hind paw and contralateral fore paw are simultaneously in stance or swing phase. Trot. |
| | **Left/Right Coupling (hind, fore)** | Left-right alternation rhythm. Ratio of time difference between consecutive left and right ground contacts to whole stride duration. |
| | **L/R Coupling Deviation (hind, fore)** | Deviation of Left/Right Coupling between the strides. |
| | **Step Width (hind, fore)** | The distance between left and right hind/fore paw during stance phase, perpendicular to midline. |
| | **Step Width Deviation (hind, fore)** | The deviation of Step Width between the strides. |
| | **Double Support (hind, fore)** | Percentage of stride time the both left and right (hind or fore) limbs simultaneously are in ground contact. |
| | **Single Support (hind, fore)** | Percentage of stride time when one limb of the hind/fore limb pair is in ground contact and the other is not. |
| | **Support Zero** | Percentage of stride time none of the four limbs are in ground contact (and all four limbs in mid air) |
| | **Support Single** | Percentage of stride time one of the four limbs is in ground contact (three in mid air) |
| | **Support Diagonal/Girdle/Lateral** | Percentage of stride time two of the four limbs are in ground contact, three modes: diagonal, girdle (galloping), lateral |
| | **Support Three** | Percentage of stride time three of the four limbs are in ground contact (one in mid air) |
| | **Support Four** | Percentage of stride time all the four limbs are in ground contact. |
| | **Toe Clearance (hind, fore)** | Maximum clearance, i.e., distance from the ground, of a paw during swing phase. |
| **Body Posture** | **Iliac Crest Height** | Height of iliac crest during mid-stance. |
| | **Mean Hip Height** | Average height of hip over a stride. |
| | **Hip Height Range** | Range of hip height (vertical movement) during a stride. |
| | **Mean Hip Jerk** | The average degree of non-smoothness, i.e., rate of acceleration change, of hip during stride. |
| | **Tail Base Height (min, mean, max)** | Minimum, average, and maximum height of tail tip from the ground. |
| | **Tail Base Height Range** | Range of vertical tail base movement during a stride. |
| | **Protraction (hind)** | Maximum protraction of hind paw with respect to iliac crest point (forward direction, occurs at initial contact) |
| | **Retraction (hind)** | Maximum retraction of hind paw with respect to iliac crest point (backward direction, occurs at initial swing) |
| | **Nose Height** | Average height of nose. |

| | | |
|---|---|---|
| | **Nose Height Range** | Range of nose height during a stride. |
| | **Lateral Head Rotation** | Average absolute value of lateral head rotation angle, based on head direction with respect to central line in horizontal plane. |
| | **Head Rotation Deviation** | Deviation of Lateral Head Rotation between different strides. |
| | **Head Rotation Range** | Range of Lateral Head Rotation angle in horizontal plane during a stride. |
| | **Tail Tip Height (min, mean, max)** | Minimum, average, and maximum height of tail tip from the ground. |
| **Tail Tip** | **Tail Tip Height Range** | Range of vertical tail tip movement during a stride. |
| | **Tail Tip Over Hip** | Percentage of stride duration the tail tip is higher than hip level. |
| | **Tail Tip Ground Contact** | Percentage of stride duration the tail tip touches ground. |
| | **Tail Tip Distance 2D** | Ratio of two-dimensional tail tip trajectory length to stride length, determined from the side view. |
| | **Tail Tip Distance 3D** | Ratio of three-dimensional tail tip trajectory length to stride length. |
| | **Hip Angle (min, mean, max)** | Hip joint angle, minimum, mean, and maximum values. |
| **Joint Angles** | **Knee Angle (min, mean, max)** | Knee joint angle, minimum, mean, and maximum values. |
| | **Ankle Angle (min, mean, max)** | Ankle joint angle, minimum, mean, and maximum values. |
| | **Hip ROM** | Hip joint range of motion (ROM) during a stride, difference between max and min Hip Angles |
| | **Knee ROM** | Knee joint range of motion during a stride. |
| | **Ankle ROM** | Ankle joint range of motion during a stride. |
| | **Hip ROM Deviation** | Deviation of hip ROM between different strides. |
| | **Knee ROM Deviation** | Deviation of knee ROM between different strides. |
| | **Ankle ROM Deviation** | Deviation of ankle ROM between different strides. |
| | **Paw Trajectory Shape 25% (hind, fore)** | Percentage of swing phase duration the paw is above 25% of Toe Clearance. |
| **Paw Trajectory** | **Paw Trajectory Shape 50% (hind, fore)** | Percentage of swing phase duration the paw is above 50% of Toe Clearance. |
| | **Paw Trajectory Shape 75% (hind, fore)** | Percentage of swing phase duration the paw is above 75% of Toe Clearance. |
| | **Toe Lift-Off Angle (fore, hind)** | Angle of paw trajectory ascent at the early swing phase. |
| | **Relative Trajectory Length** | Ratio of fore paw 2D trajectory path length to stride length, subtracted by one. |
| | **Excess Vertical Movement** | Ratio of vertical fore paw trajectory distance to double of Toe Clearance, subtracted by one. |
| | **Backward Paw Distance** | Sum of excess backward movement of forepaw during a stride. |

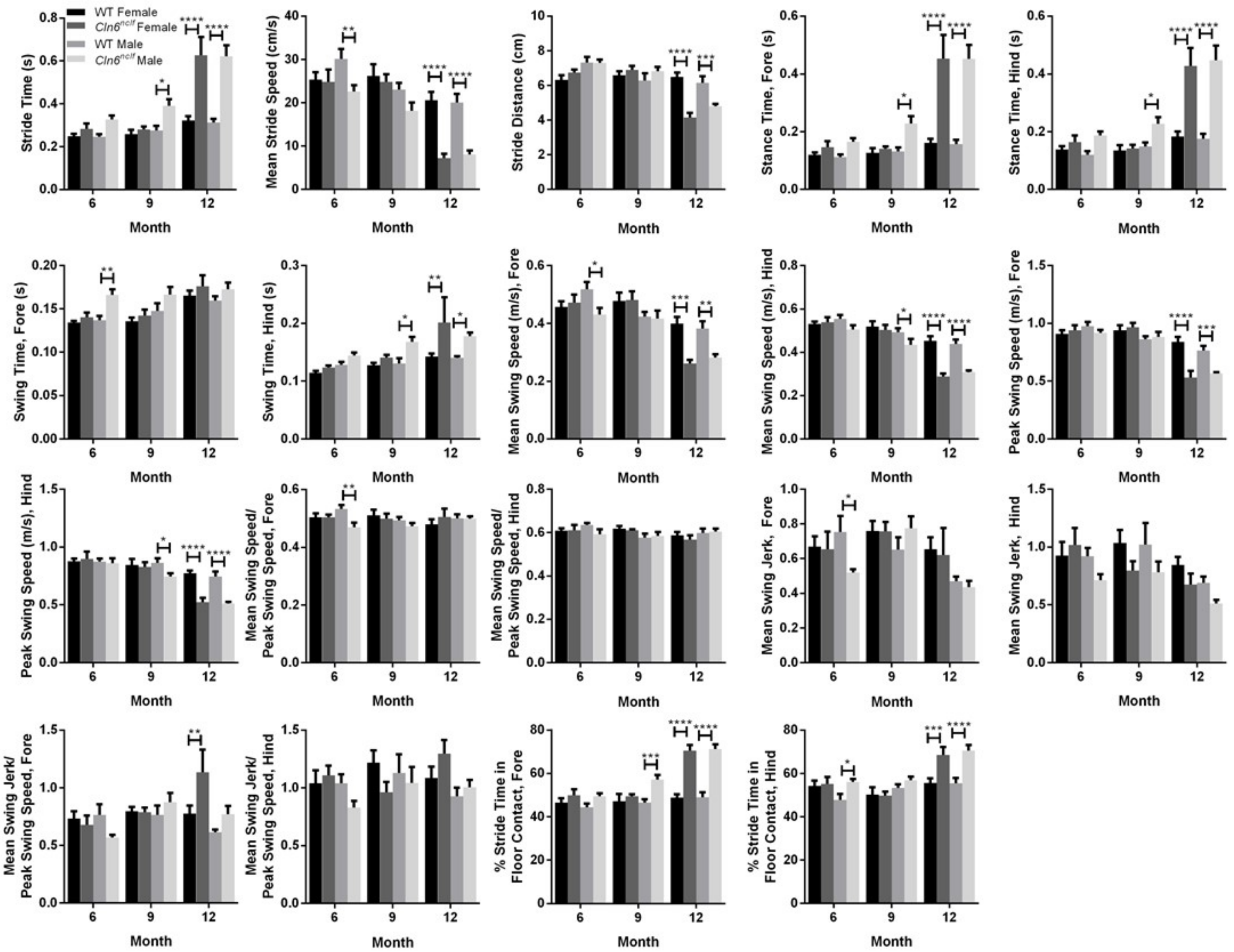

**Fig. S4: Spatiotemporal kinematic parameters**. WT and *Cln6$^{nclf}$* mice were observed from 6-12 months period. Data is mean ± SEM, n = 12 for WT (6 male, 6 female; data pooled), n = 11 for *Cln6$^{nclf}$* (5 male, 6 female). Statistical significances: unpaired t-test **p* < 0.05, ***p* < 0.01, ****p* < 0.001, *****p* < 0.0001.

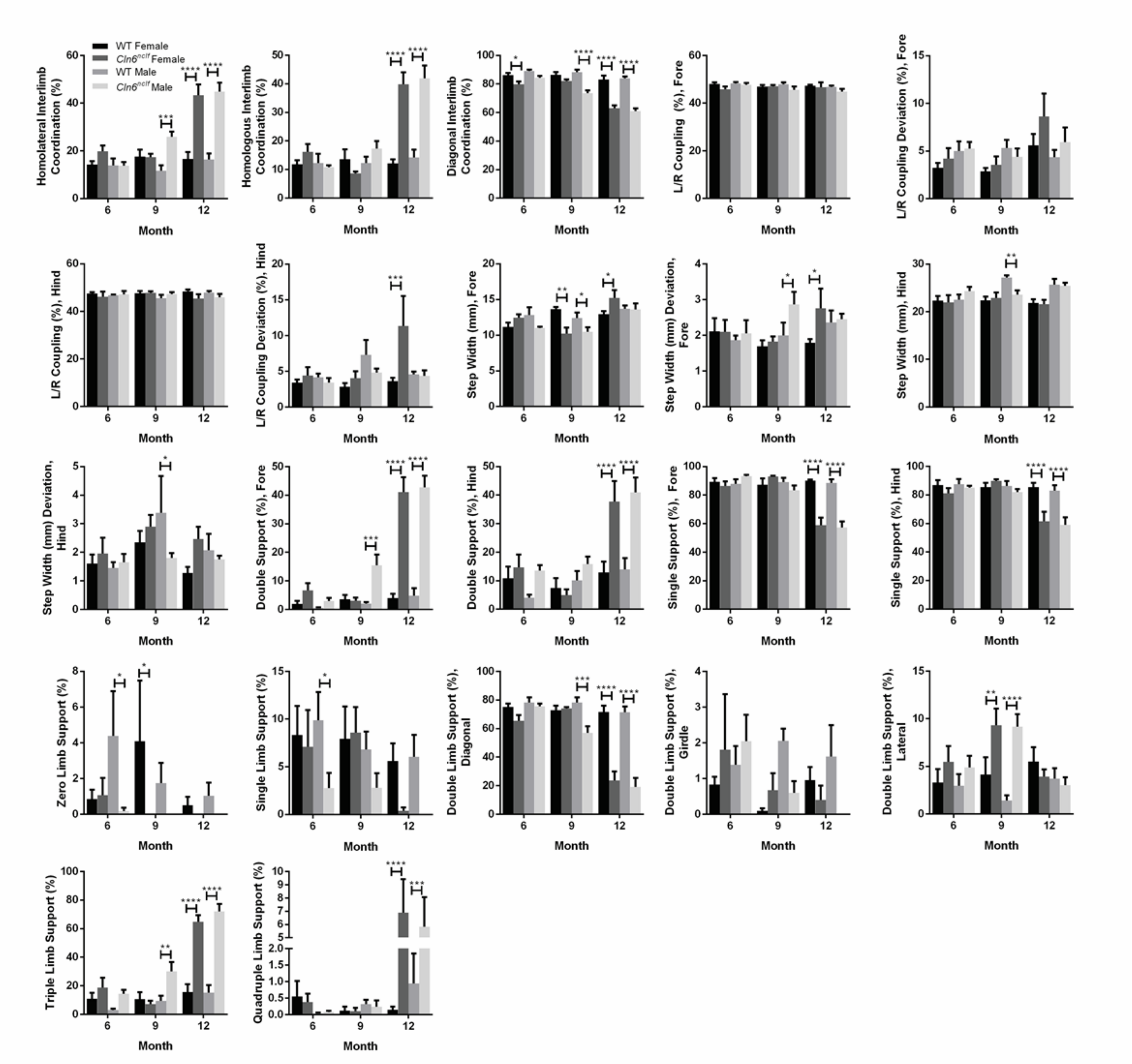

**Fig. S5: Interlimb coordination parameters.** WT and *Cln6^nclf* mice were observed over 6-12 months period. Data is mean ± SEM, n = 12 for WT (6 male, 6 female; data pooled), n = 11 for *Cln6^nclf* (5 male, 6 female). Statistical significances: unpaired t-test *p < 0.05, **p < 0.01, ***p < 0.001, ****p < 0.0001.

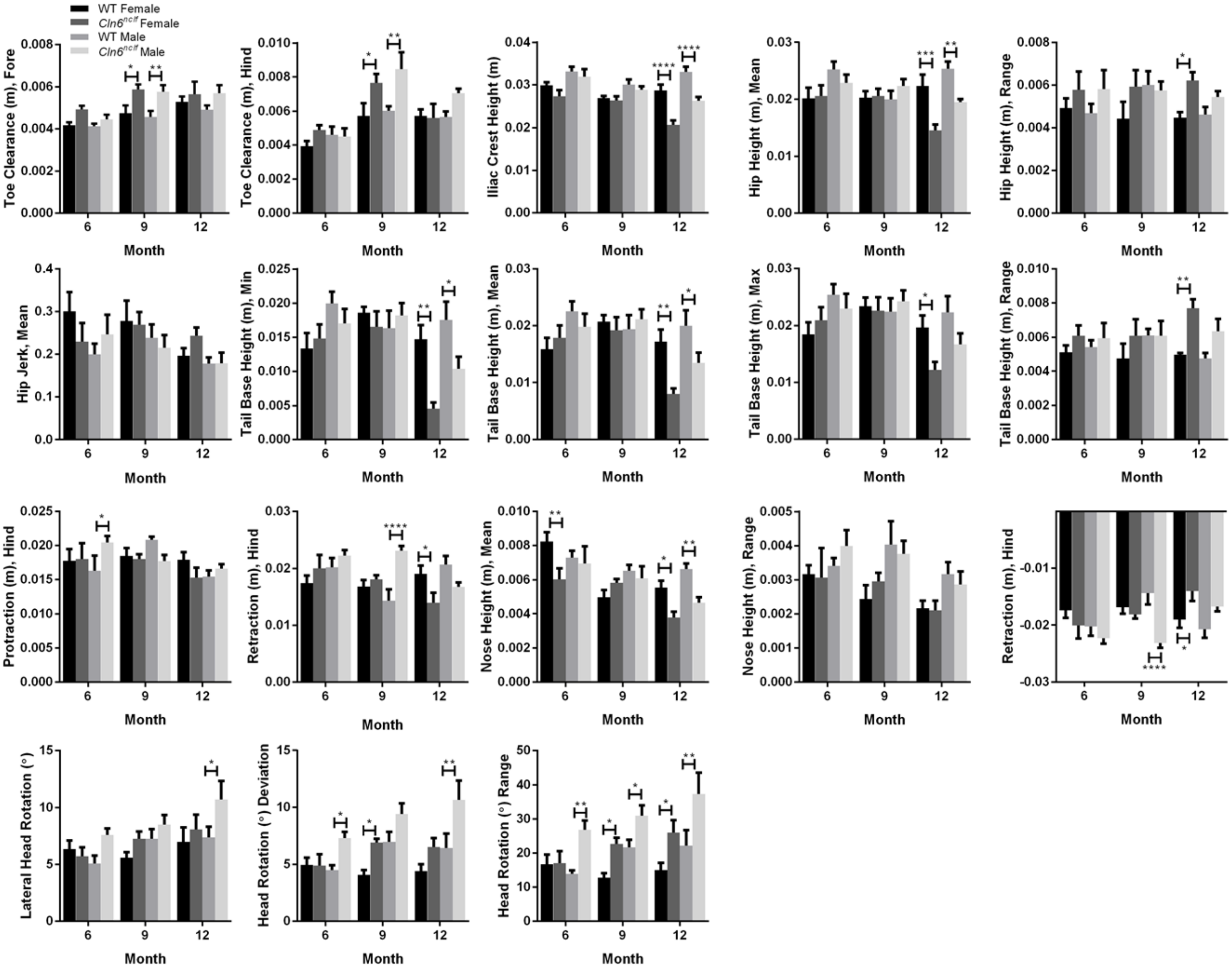

**Fig. S6: Kinematic parameters describing body posture, toe clearance, hind limb protraction and retraction, nose height and head rotation**. WT and *Cln6$^{nclf}$* mice were observed over 6-12 months period. Data is mean ± SEM, n = 12 for WT (6 male, 6 female; data pooled), n = 11 for *Cln6$^{nclf}$* (5 male, 6 female). Statistical significances: unpaired t-test *p < 0.05, **p < 0.01, ***p < 0.001, ****p < 0.0001.

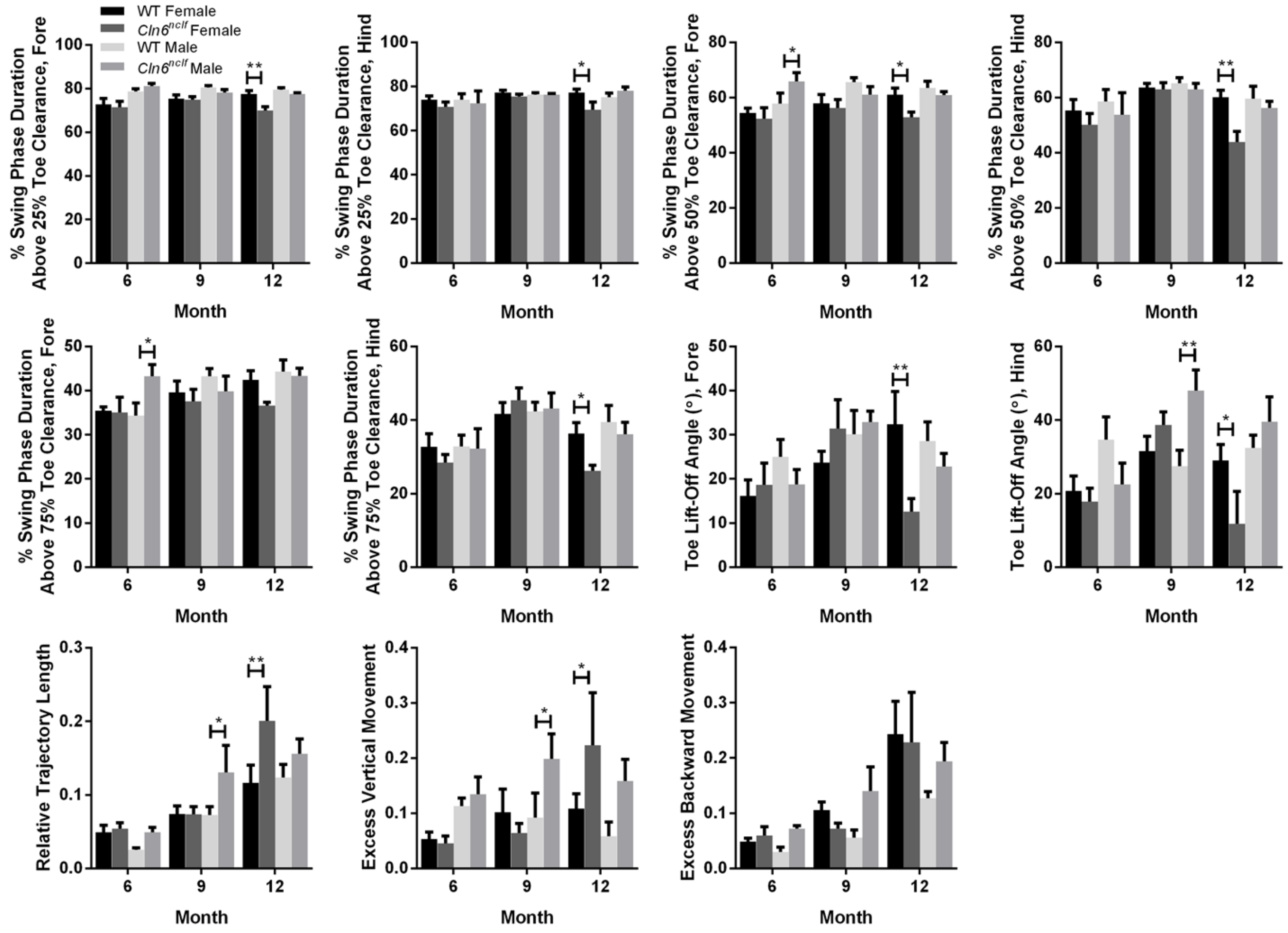

Fig. S7: **Kinematic parameters describing limb trajectory profiles and excess movements during swing phase of gait**. WT and *CLN6*$^{nclf}$ mice were observed over 6-12 months period. Data is mean ± SEM, n = 12 for WT (6 male, 6 female; data pooled), n = 11 for *CLN6*$^{nclf}$ (5 male, 6 female). Statistical significances: unpaired t-test *$p < 0.05$, **$p < 0.01$, ***$p < 0.001$, ****$p < 0.0001$.

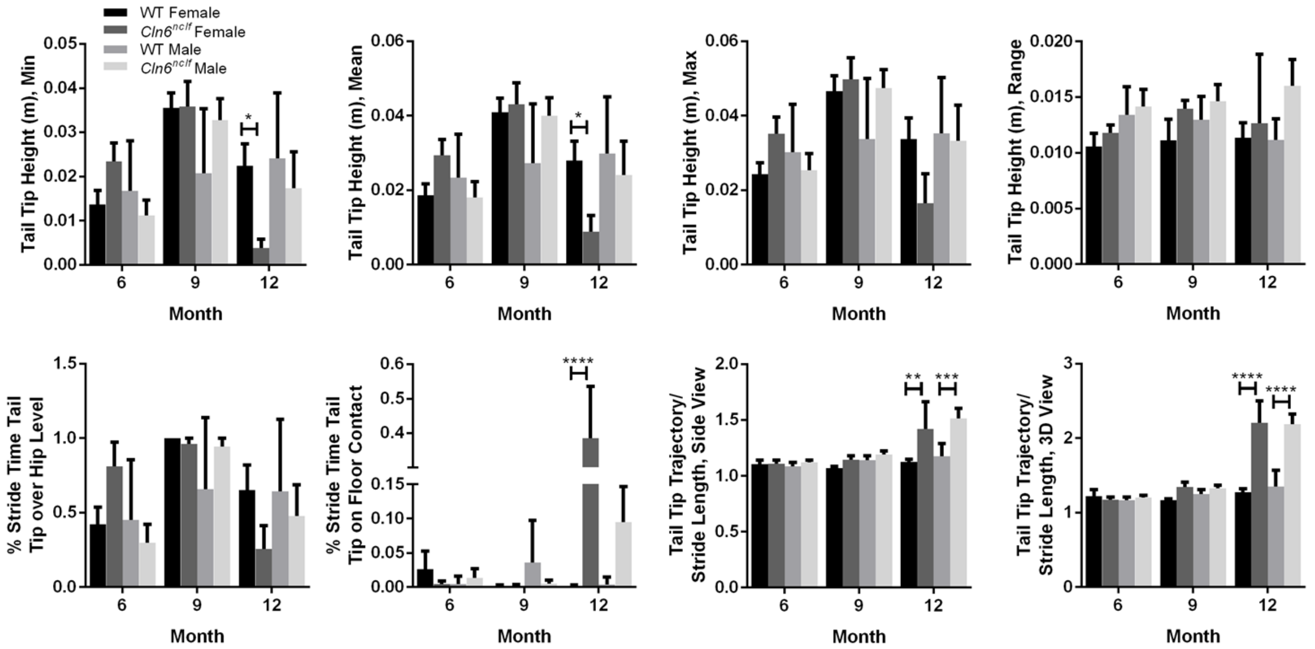

Fig. S8: **Kinematic parameters describing tail tip movements**. WT and *Cln6*$^{nclf}$ mice were observed over 6-12 months period. Data is mean ± SEM, n = 12 for WT (6 male, 6 female; data pooled), n = 11 for *Cln6*$^{nclf}$ (5 male, 6 female). Statistical significances: unpaired t-test *$p < 0.05$, **$p < 0.01$, ***$p < 0.001$, ****$p < 0.0001$.

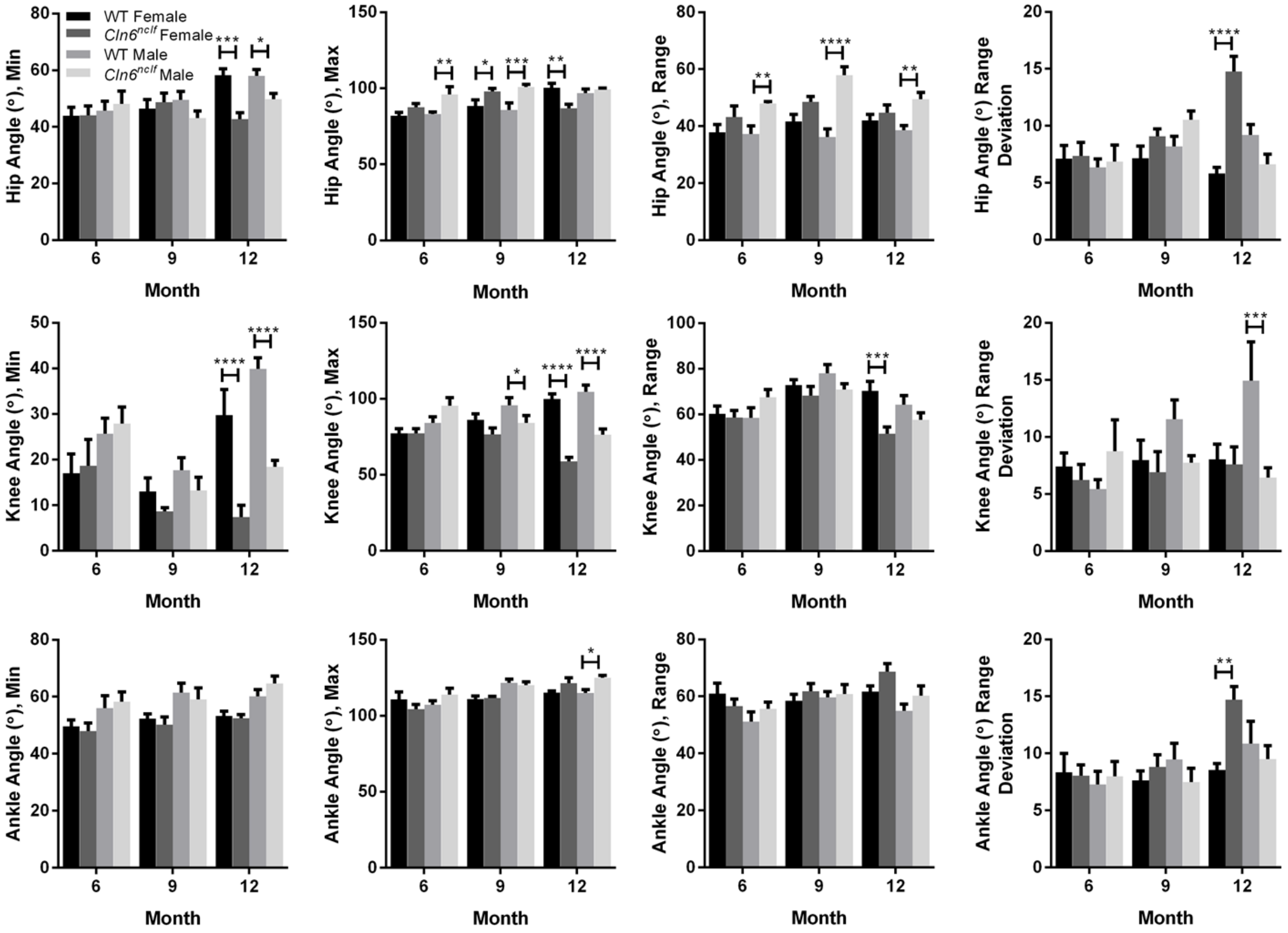

**Fig. S9: Hip, knee and ankle angles.** WT and *Cln6*$^{nclf}$ mice were observed over 6-12 months period. Data is mean ± SEM, n = 12 for WT (6 male, 6 female; data pooled), n = 11 for *Cln6*$^{nclf}$ (5 male, 6 female). Statistical significances: unpaired t-test *$p < 0.05$, **$p < 0.01$, ***$p < 0.001$, ****$p < 0.0001$.
